# Simulating Multiple Substrate Binding Events by γ-Glutamyltransferase using Accelerated Molecular Dynamics

**DOI:** 10.1101/2020.04.20.050500

**Authors:** Francesco Oliva, Jose C. Flores-Canales, Stefano Pieraccini, Carlo F. Morelli, Maurizio Sironi, Birgit Schiøtt

**Author notes:** **Corresponding Author** Stefano Pieraccini –, Birgit Schiøtt –. **Author Contributions** F.O. and J.C.F.-C. have contributed equally to this work.

## Abstract

γ-glutamyltransferase (GGT) is an enzyme that uses γ-glutamyl compounds as substrate and catalyzes their transfer into a water molecule or an acceptor substrate with varied physiological-function in bacteria, plants and animals. Crystal structures of GGT are known for different species and in different states of the chemical reaction; however, structural dynamics of the substrate binding to the catalytic site of GGT is unknown. Here, we modeled *Escherichia Coli* GGT’s glutamine binding by using a swarm of accelerated molecular dynamics (aMD) simulations. Characterization of multiple binding events identified three structural binding motifs composed of polar residues in the binding pocket that govern glutamine binding into the active site. Simulated open and closed conformations of a lid-loop protecting the binding cavity suggests its role as a gating element by allowing or blocking substrates entry into the binding pocket. Partially open states of the lid-loop are accessible within thermal fluctuations, while the estimated free energy cost of a complete open state is 2.4 kcal/mol. Our results suggest that both specific electrostatic interactions and GGT conformational dynamics dictate the molecular recognition of substrate-GGT complexes.

## Introduction

Molecular recognition of substrate molecules by enzymes is characterized by substrate binding to a catalytic site. Upon binding, the protein explores the conformational space to find a transition state lowering the activation barrier of a chemical reaction followed by a product release stage^1^. The timescale of this process ranges from microseconds to milliseconds^2–4^. Early during a substrate recognition, the ligand samples rotational, translational and conformational degrees of freedom in order to reach a bound state over several microseconds^5,6^. The substrate binding process is critical for enzyme activity and specificity. Hence, an atomistic description of the ligand entry and binding into the catalytic site and the protein conformational space explored upon binding is highly desirable both to better understand the origin of such specificity and to pursue the rational design of improved enzymes for chemical synthesis^7^. In this work, we focused our attention on γ-glutamyltransferase, also referred to as γ-glutamyltranspeptidase (GGT, E.C. 2.3.2.2), a heterodimeric enzyme able to cleave amide bonds and widely distributed from bacteria to plants and mammals^7,8^. Physiological importance of GGT is also diverse, extending from a primary role in maintaining cysteine levels in animal and plant cells^9–12^, nitrogen stress response in yeast cells^13^, to a virulence factor of *Helicobacter pylori*, a bacteria, that enables its widespread infection responsible for chronic gastritis and increased risk of gastric cancer^14^.

All the members of the GGT superfamily are autocatalytically activated through a proteolytic cleavage, which liberates a N-terminal nucleophilic, catalytically active residue^15^. GGT cleaves γ-glutamyl compounds, e.g. glutathione, releasing the cysteinylglycine portion and forming a covalently bound γ-glutamyl-enzyme intermediate through formation of an ester linkage between the γ-carbon atom of the glutamyl group and the oxygen atom of the catalytically active, N-terminal Thr391 of the small enzyme subunit^16^ (Figure 1a). The γ-glutamyl-enzyme intermediate then either reacts with a water molecule and undergoes hydrolysis or it transfers the glutamyl group to an amino acid or a short peptide (transpeptidation)^8,17–19^ (Figure 1b).

**Figure 1.**
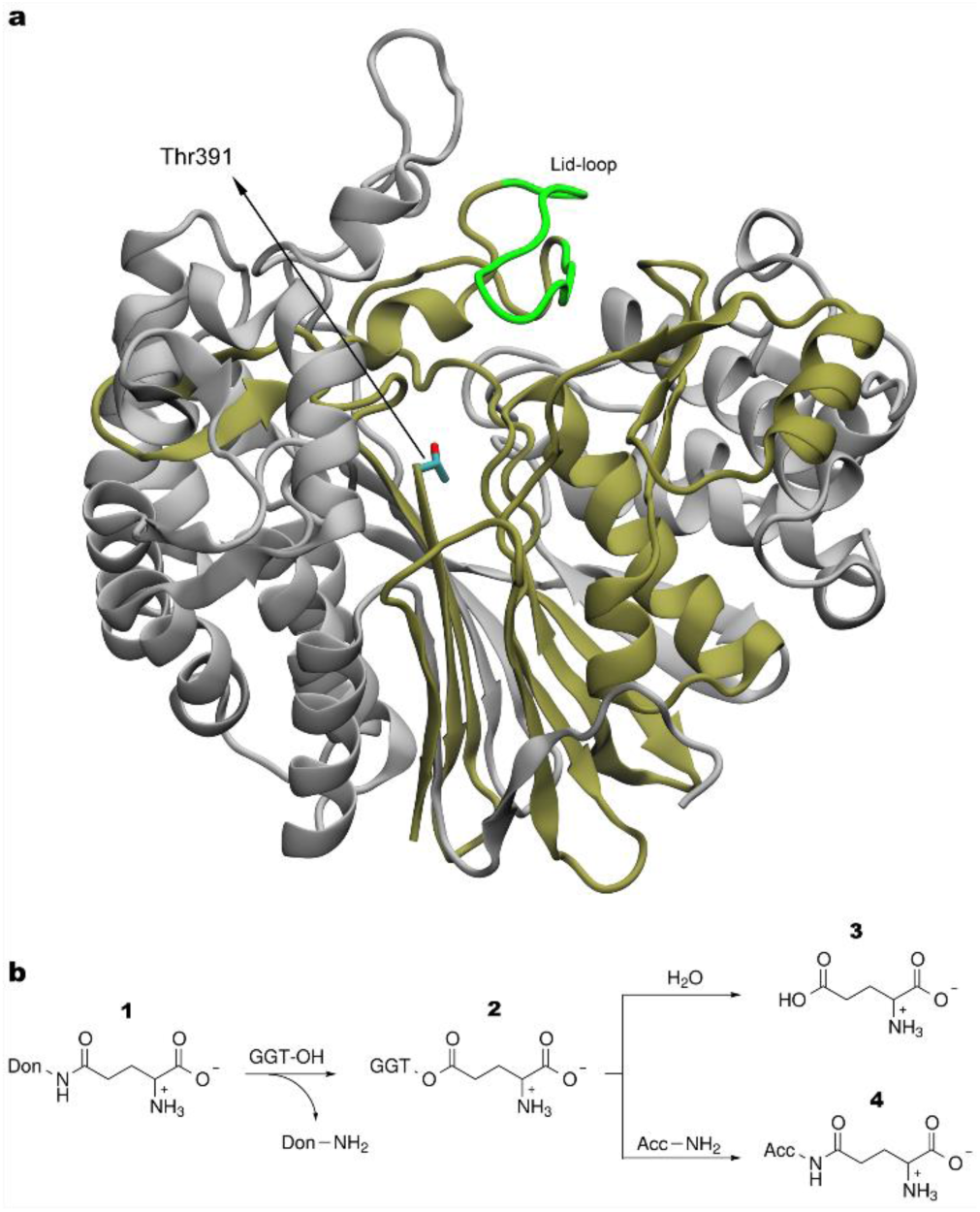
Structure representation of *Escherichia Coli* GGT (EcoGGT) and catalytic reaction. **a**, X-ray structure of EcoGGT (PDB ID code 2DBX). Grey and tan color ribbons represent the heavy and light subunits, respectively. Nitrogen, carbon and oxygen atoms are represented in blue, cyan and red color, respectively. **b**, Schematic representation of GGTs mechanism. Reaction between GGT and the γ-glutamyl donor **1**, formation of the γ-glutamyl enzyme intermediate **2**. The intermediate can react in two ways: with water resulting in hydrolysis and liberation of glutamic acid **3**; or with a peptide resulting in a new γ-glutamyl derivative **4** through a transpeptidation reaction.

Here, we focus on *Escherichia Coli* GGT (EcoGGT). Its catalytic cycle has been characterized and it is known to be localized in the periplasm; however, its physiological role is unknown^20^. EcoGGT found use also as a biocatalyst for the synthesis of γ-glutamyl derivatives at a preparative level^21^, accepting L-but not D-amino acids and has a preference for basic and aromatic acceptor amino acids^22^. This contrasts with its low stereo-specificity for γ-glutamyl substrates. Crystal structures of EcoGGT have been solved in the apo state, in complex with an intermediate and the corresponding hydrolysis product^23,24^ and, with the aid of structural information, light was shed on the autocatalytic process leading to enzyme maturation^25^ and on the reaction mechanism^26^. Based on these structures, a so-called lid-loop (residues 438 to 449) was found to form a hydrogen bond between residues Asn411 – Tyr444 closing the entry to the binding pocket. This lid-loop was also found to be conserved among other bacterial and human GGT sequences^27^. It has been pointed out that the observed closed lid-loop conformations in crystal structures of *E*. *Colit* and *Helicobacter pylori* GGTs contribute to their lower activity compared to human GGT^24,27^. The crystal structure of the human GGT revealed an open lid-loop providing larger access to the active site. Despite the research progress in understanding the catalytic stages of GGT enzymes, no structural and dynamical information of the substrate binding and the role of the lid-loop in GGTs has been obtained. Elucidation of the binding process will provide insight on the physicochemical principles dictating the activity and specificity of the GGT family.

Molecular dynamics (MD) simulations are increasingly used to get insight into fundamental biomolecular processes such as protein folding, allostery regulation, and protein-ligand binding^28–30^. Conventional MD simulations are usually limited to a few microseconds in conventional high-performance computing clusters, which limit the modeling of multi-microsecond time scale processes such as protein-ligand binding. To overcome these limitations, several enhanced sampling methods, such as replica exchange MD^31^, metadynamics^32^, umbrella sampling^33^, transition path sampling^34^ and accelerated molecular dynamics (aMD)^35^, have been developed to speed up phase space exploration. In particular, aMD is an enhanced sampling technique that works by reducing energy barriers of the molecular potential energy surface, adding a non-negative boost potential when the potential energy of the system is lower than a reference energy^35^. It has been employed to simulate rare events that occurs on the microseconds to milliseconds time scale, such as protein folding and ligand-protein binding, with tens to hundreds of nanoseconds long simulations^36–41^.

In this work, we performed an ensemble of short aMD repeats and focused on a subset of trajectories where GGT-glutamine binding was observed. Our approach allowed us to model the process of substrate binding by the enzyme and to identify polar residues in the binding pocket governing glutamine binding, from the free unbound state to its full insertion into the active site. The EcoGGT’s lid-loop was also found to function as a gating component during the molecular recognition process. Notably, no restraint was applied and no prior information or hypothesis concerning the enzyme-substrate interactions were used in the simulations.

## Methods

### Accelerated Molecular Dynamics Simulations

Molecular recognition of glutamine by EcoGGT and the subsequent formation of the ligand-protein complex was simulated with aMD. This method was first introduced by McCammon et al^35^ and designed to accelerate transition events between different conformational minima. This is obtained by boosting the potential energy every time the value of the true potential is below a fixed threshold.

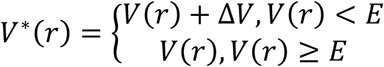

Where *V*^*^(*r*) is the modified potential, V(r) is the real potential and E is the energy threshold. The implementation of the method that we adopted is the “dual boost” approach^42^. With this method, two separated boosts are applied, one to the torsional terms and a separate one to the total potential energy.

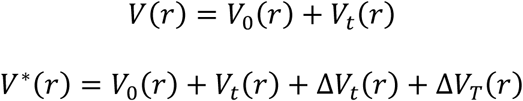

Where *V*_*t*_(*r*) is the total potential of the torsional terms, Δ*V*_*t*_(*r*) and Δ*V*_*T*_(*r*) are the boost potentials applied to the torsional term *V*_*t*_(*r*) and the total potential, respectively. This approach allows to accelerate rare events and increase the diffusion without destroying the local water structure^42^. A more detailed explanation of the dual boost method can be found in the Supporting Material (SM).

### System Preparation

The structure of EcoGGT was obtained from the protein data bank (PDB ID code 2DBX^24^). The crystal structure contains a dimer of the protein and the subunits are labeled A, B, C and D. The resolution was 1.7 Å. Subunits C and D were removed as well as all the water molecules, two calcium ions, one glycerol and one glutamate molecule. In the starting structure of each aMD run, glutamine was placed 25 Å away (distance between the hydroxyl oxygen of Thr391 and glutamine carboxyl carbon), from EcoGGT and each run has different random seed numbers and initial velocities. The system was simulated in explicit water solvent employing periodic boundary conditions with initial box dimensions 100 Å × 101 Å × 92 Å.

### Simulation Details

Glutamine and EcoGGT were described with the amber99SB^43^ force field and the TIP3P^44^ model was adopted for water. The total number of atoms was 89736, divided in 8093 protein atoms, 20 glutamine atoms, 14 sodium counter-ions and 81608 water atoms (27203 water molecules). We optimized the initial system geometry with a 50000 steps of energy minimization, using OpenMM 7.2.1 CUDA kernels^45^. All covalent bonds to hydrogens were constrained to their equilibrium values with the SETTLE algorithm^46^. Position restraints were imposed on heavy atoms in the protein with a spring force of 100 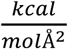. We then carried out a 40 ps equilibration in the NVT ensemble, slowly increasing the temperature from 50 to 300 K in 4 steps and progressively relaxing position restraints from 100 to 4 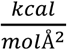 in 5 steps. To control the temperature we used a Langevin integrator^47^. A time step of 2 fs was used for all the reported MD runs. The NVT equilibration simulation stage was followed by a second equilibration of 50 ps in the NPT ensemble, during which restraints were fully removed in 5 steps. Pressure control was assured through an isotropic Monte Carlo barostat which attempts to change the volume every 25 steps and accepts the changes with the Metropolis Monte-Carlo criterion^48,49^. This was followed by an equilibration run of 10 ns and a production run of 50 ns. aMD parameters were calculated using the respective total and dihedral averages from the 50 ns production run. Table S1 provides the values used to calculate aMD parameters, further described in the Supplemental Material. aMD protocol implementation was based on a dual-boosting aMD python script available at the OpenMM virtual repository^50^.

### Analysis Details

Phrases clustering analysis: This is a clustering method for the identification of clusters of protein residues making simultaneous contacts with a ligand through an MD trajectory^51,52^. It uses a distance metric based on the surprisal of a pair of residues to be in contact with a ligand: *d*(*i, j*) = −*log* ((*n*(*i, j*) + 1)/*N*), where n(i, j) is the number of simultaneous contacts for residues i and j over a trajectory containing N frames. Note that *d*(*i, j*) ∼ 0 and *d*(*i, j*) = *log* (*N*) correspond to residues i and j always in contact with the ligand, and no simultaneous contacts throughout the trajectory, respectively. A ligand and a protein residue are in contact if any of their heavy-atoms are within 3 Å of each other. Clustering is carried out using complete-linkage hierarchical clustering. A heatmap plot is used to summarize the data and is truncated to values lower than 4 to focus on specific protein-ligand interactions. The analysis was carried out using MD frames where the ligand is bound to the protein, see below. The source code is available at https://gitlab.au.dk/Biomodelling/Phrases.git.

Clustering analysis: Glutamine-enzyme complexes are extracted from the trajectories where a binding event occurs (runs 10, 20 and 50 from now on referred to as R10, R20 and R50, respectively), and concatenated into a single data set when the root-mean-square deviation (RMSD) value between glutamine heavy-atoms and a model of the enzyme bound to glutamine (further described in the Results section), is steadily below 10 Å. The binding pocket flexibility is studied by clustering analysis performed^53^ using the heavy-atoms of glutamine, residues in the binding pocket (Ser462, Asn411, Ser463, Gly484, Met464, Gly483, Asp433, Arg114, Gln430, Thr409, Thr391, Ser485), and residues undergoing conformational changes Pro482 and Tyr444. The conformational space sampled by glutamine bound in the binding pocket was analyzed by clustering on a substrate heavy-atom RMSD basis. The GROMOS clustering method was employed for all reported clustering analysis using a 1 Å cut-off distance^54^. The central structure of each cluster is defined as the structure with the largest number of neighbors. Protein graphs rendering was performed with VMD^55^. Two dimensional (2D) plots of the substrate-protein interactions were obtained with Ligplot+^56^. A VMD script used to calculate the number of water molecules between Tyr444 and the substrate is included in the Supplemental Material.

Lid-loop Free Energy Profile: The accumulated data of all aMD trajectories was binned into 25 bins such that the minimum number of aMD frames in a bin is 10. The profile was estimated by direct reweighting of each aMD snapshoot and the bias exponential factors were approximated using a Maclaurin series expansion of order 10^57^.

## Results

The starting simulation system consists of the EcoGGT heterodimer structure in the presence of one unbound glutamine, while solvated in explicit solvent. We performed 60 aMD independent runs, each of them with a simulation length of ca. 100 ns, in high-throughput computing clusters.

### Glutamine Binding to the EcoGGT Catalytic Site

As the nucleophilic attack of the hydroxyl oxygen of Thr391 on the glutamine amide carbon is the first step in the formation of the γ-glutamyl-enzyme intermediate, this C-O distance has been monitored along the trajectories, in order to select those leading to potentially reactive bound conformations. This is in line with the near-attack conformation (NAC) theory where a substrate adopts a ground-state conformation, defined by the distance between reactive atoms and/or angles, before bond making and breaking^58^. In 5 out of 60 trajectories the C-O distance fell below 10 Å, showing that the substrate approached the protein active site. In 3 of these 5 trajectories, the substrate durably binds into the catalytic site until the end of the MD trajectories (trajectories named R10, R20 and R50 in Figure 2a). Whereas, in the other two trajectories, namely R22 and R30, glutamine rapidly escaped from the binding pocket region after less than 20 ns (Figure 2a). Trajectories R20 and R50 show similar final values, while R10 stabilizes around a larger nucleophilic attack distance indicating a different substrate-enzyme pocket conformation. This will be further analyzed in a following subsection. The small number of substrate-enzyme binding observations in an ensemble of short aMD runs mirrors the behavior of real-world enzymes, where only a small fraction of enzyme-substrate collisions has the proper steric and energetic requirements to evolve into a reactive encounter.

**Figure 2.**
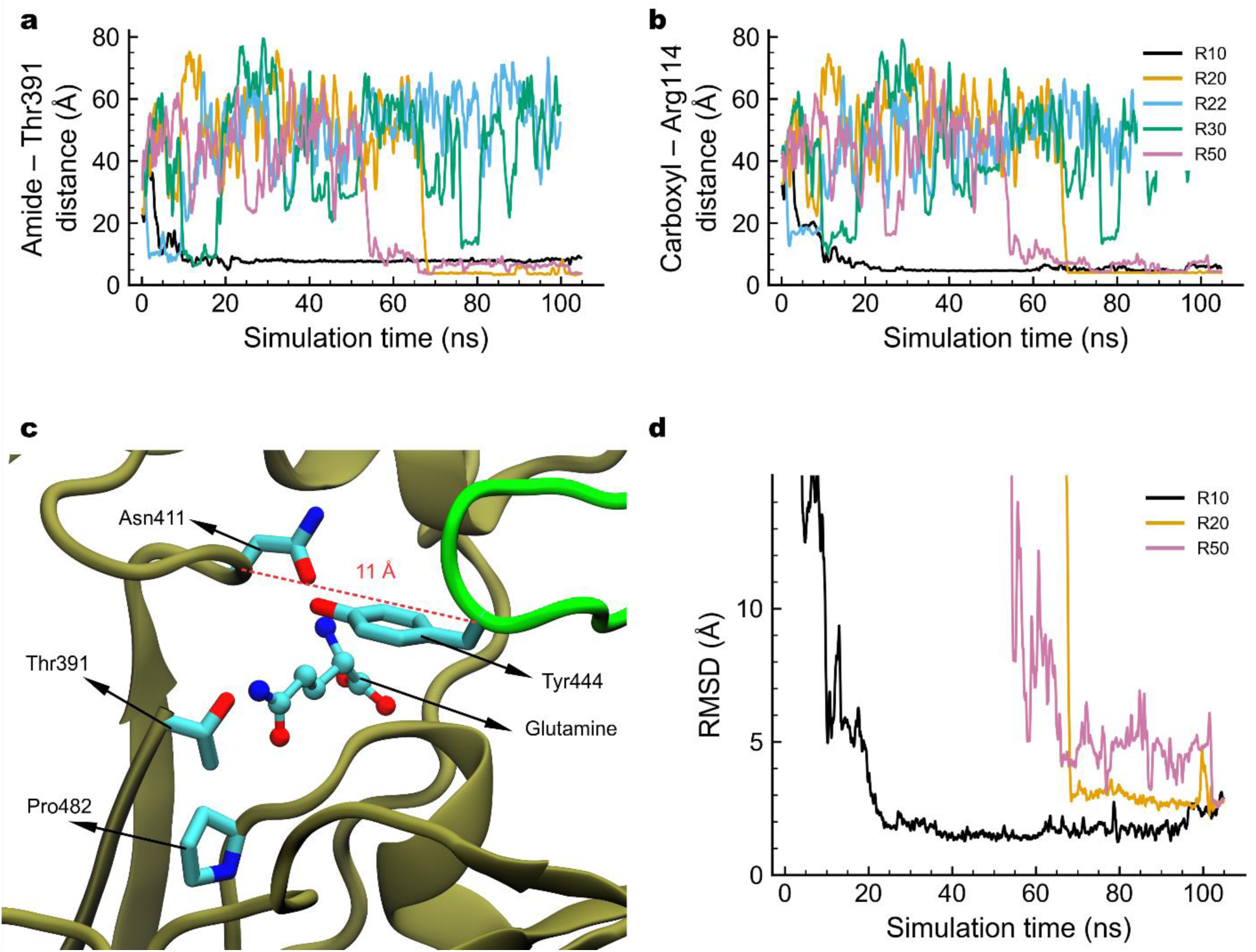
Substrate binding by EcoGGT. **a**, Distance between the glutamine amide carbon and the reactive hydroxyl oxygen atoms on Thr391. **b**, Distance between the glutamine carboxyl carbon and the guanidine carbon in Arg114. Trajectories named R10, R20, R22, R30 and R50 are shown. **c**, Glutamine-EcoGGT complex model built from a cocrystallized glutamate-enzyme structure. Lid-loop and chain B of EcoGGT are shown in green and tan color ribbons, respectively. Cα distance between Asn411 and Tyr444 is highlighted. Nitrogen, carbon and oxygen atoms are represented in blue, cyan and red color spheres, respectively. **d**, RMSD traces of glutamine with respect to the enzyme-glutamine complex model. Trajectories were aligned with respect to the crystal X-ray structure and the RMSD values were calculated using only the substrate heavy atoms. Data series are smoothed by a moving average over 20 frames (400 ps).

Comparison of the five trajectories shows that a salt bridge interaction between glutamine C-terminus and Arg114 is essential for a stable binding of glutamine into the binding pocket. The distance between the glutamine carboxyl carbon and the Arg114 guanidinic carbon (Figure 2b) steadily falls below 5 Å in successful trajectories. The abovementioned five trajectories are reported in detail in the rest of this article.

### Comparison of Simulated Glutamine Binding to the EcoGGT-Glutamate Complex

Simulated EcoGGT-substrate bound structures are compared to EcoGGT co-crystallized with glutamate, the product of the enzymatic reaction as shown in the reaction scheme in Figure 1b^24^. Due to the size and shape similarity between glutamine and glutamate, it is hypothesized that the position and orientation of the two molecules are similar before and after the reaction takes place (Figure 2c). This is tested by aligning the aMD generated protein structures to the atomic coordinates of the crystal structure (PDB ID 2DBX), followed by RMSD calculations using the substrate heavy-atoms (Figure 2d).

Figure 2d shows that the substrate samples bound structures similar to a manually constructed model using the crystallographic structure of the product and chemical intuition. In R10, glutamine spends more than 80 ns with an average RMSD of ca. 2 Å relative to the proposed substrate bound conformation, while in R20 and R50 the substrate samples conformations with RMSD values ca. 3 Å over the last 30 and 3 ns, respectively.

### Specific Glutamine Binding Site Interactions

Structural binding motifs are identified in our aMD simulations by identifying pairs of residues simultaneously in contact with the substrate followed by hierarchical clustering (see Phrases clustering description in Methods). Phrases is based on information theory and provides a distance metric of how frequently two residues form close contacts with a ligand. The smaller the distance metric, the more frequently a specific ligand-residues interaction is sampled. Hierarchical clustering using this distance metric allows for the identification of structural motifs in ligand-protein MD simulations. Figure 3a shows a heatmap and a dendrogram plot summarizing specific substrate-pocket interactions obtained from the concatenated substrate-bound segments of trajectories R10, R20 and R50, see details in Methods. Visual inspection of the dendrogram reveals two groups of residues denominated I (Asn411, Ser462, Ser463, Met464, Gly483 and Gly484), and II (Asp433, Arg114, Gln 430, Thr409, Thr391 and Ser485). Hierarchical clustering shows a sub-cluster of 3 polar residues in group I (Asn411, Ser462 and Ser463), of which Asn411 and Ser462 frequently form ligand-coordinated contacts with residues in both groups I and II. The location of the residues in the binding pocket can explain this analysis, where Asn411 lies inside the pocket entrance close to the reactive residue Thr391 (Figures 3b-c). Meanwhile, Ser462 is located at the near opposite end of the pocket entrance allowing it to form simultaneous contacts with most residues in both groups. Group II is characterized by its highly polar nature and hierarchical clustering divides it into two motifs. One composed of residues Asp433, Arg114 and Gln430 and the other one made of residues Thr409 and Thr391. These five residues plus Asn411 are located on one side of the binding pocket opposite to the lid-loop and stabilize the substrate through electrostatic interactions. This is complemented by contact analysis calculations, which show that Asn411, Arg114, Thr409 and Asp433 form the most frequent contacts with glutamine in each of the three trajectories R10, R20 and R50 (see Contact Analysis and Figures S1-S6 in Supplementary Material).

**Figure 3.**
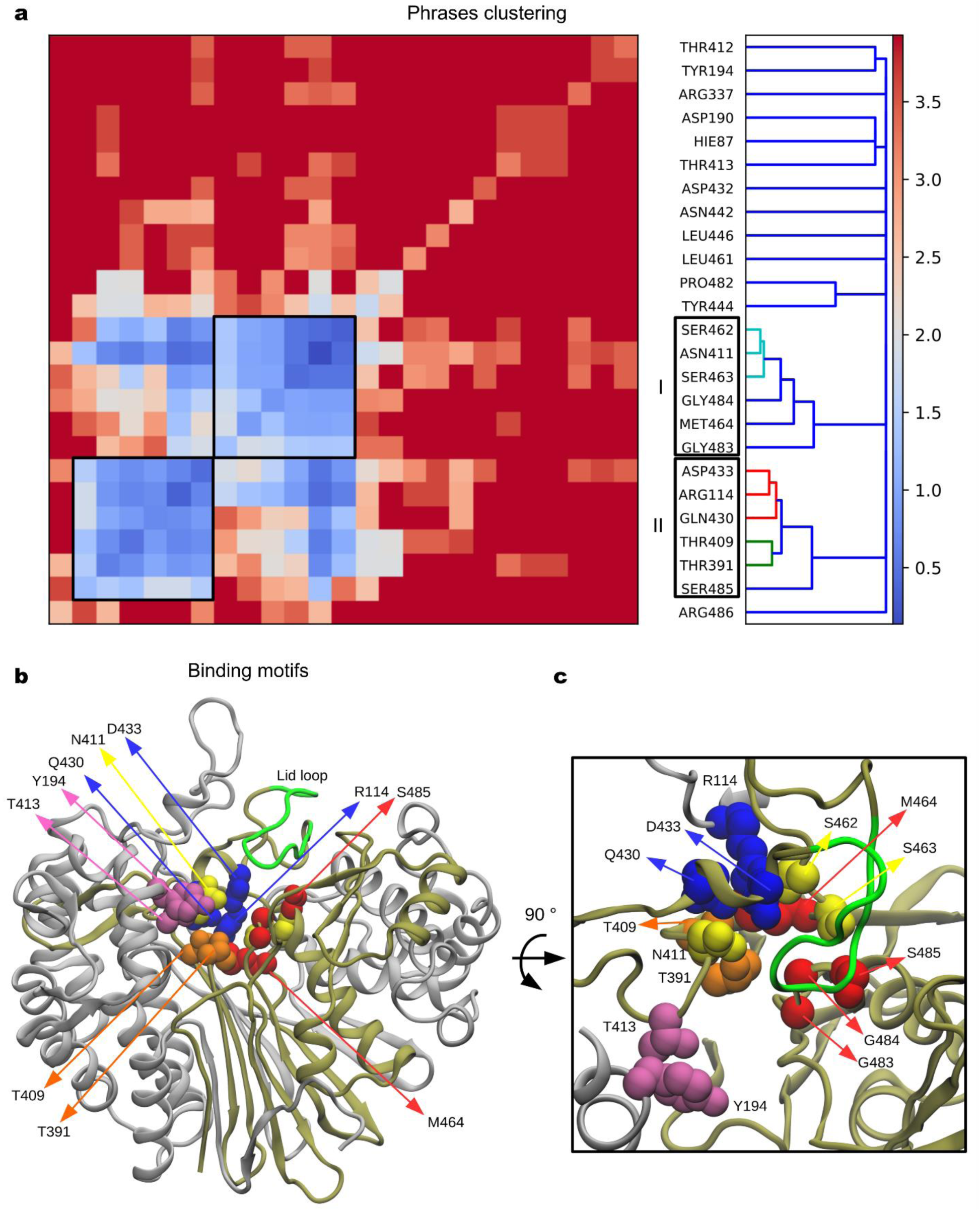
Substrate binding motifs. **a**, Hierarchical clustering of pairs of residues forming contacts with glutamine. Groups I and II are further subdivided in three binding motifs, see the dendrogram tree. **b**, Structural representation of residues forming simultaneous contacts with glutamine. Subunits A and B in silver and tan ribbons representations. Lid loop is shown in green ribbons (residues 438-449). Contacting residues are highlighted: Group I (yellow and red color sidechains in space filling representation), and group II (blue and orange color sidechains). Residues involved in the initial binding process are shown in magenta color representation. **c**, Top view after a 90 ° rotation and zooming on the access pathway.

The identification of binding motifs is coupled with further hydrogen bonding analysis for each glutamine binding trajectory (R10, R20 and R50). Our analysis suggests a common hydrogen bond pattern formation through the binding process. In all three successful trajectories, residues Thr413 and Tyr194 form short-lived hydrogen bonds with the substrate as it approaches the binding pocket (Figures 3b-c, S7). Other residues such as Thr412, Asp190 and His87 (neutral with a hydrogen on the epsilon nitrogen), are found to form short lived hydrogen bonds during the early binding process in some of the glutamine binding trajectories. Five contacting residues, of which two belong to group I (Ser463 and Asn411) and three are in group II (Arg114, Asp433 and Thr409), form hydrogen bonds when the substrate is deeply buried into the binding pocket (Figure S7). This set of five residues is hypothesized to stabilize the glutamine reactive amide carbon close to the catalytic site (Thr391). Overall, electrostatic interactions guide the substrate into the binding pocket, and phrases clustering analysis has identified three residue sub-clusters or structural motifs interacting with the substrate to fulfill this task.

### Structural Role of the Lid Loop

A loop, consisting of twelve residues (438-449), forms a lid over the substrate binding pocket, green loop in Figure 1. Note that Tyr444 on the loop tip is positioned at the pocket entrance and further forms a hydrogen bond with Asn411 in the crystal structure (Figure 2c). Lid-loop backbone conformational changes are traced through the Cα-distance between Tyr444 – Asn411 (Figure 4a). This shows that the lid-loop samples open and closed conformations in all aMD runs, whereas in the apo state the loop samples a closed conformation with short lived changes of a few nanoseconds over a 1 microsecond-long conventional MD simulation (Figure 4b). Note that R10 has the loop in a closed conformation after the substrate binding ca. 20 ns (Figure 4a). Similar lid-loop conformational changes are obtained by measuring the minimum distance between Tyr444 – and Asn411 (Figure S8). Figure 4c shows the free-energy profile of the Cα-distance between Tyr444 – Asn411, which is estimated using all 60 aMD trajectories generated in this study. Histogram of all aMD frames is shown in Figure S9. The free-energy profile has a global minimum placed at 12.6 Å and no barriers in the region between 16 to 29 Å. The region between 9 to 14 Å is within thermal fluctuations of the free energy minimum. The free energy cost of a complete open state (at 26.3 Å), of the lid-loop is ca. 2.4 kcal/mol relative to the minimum.

**Figure 4.**
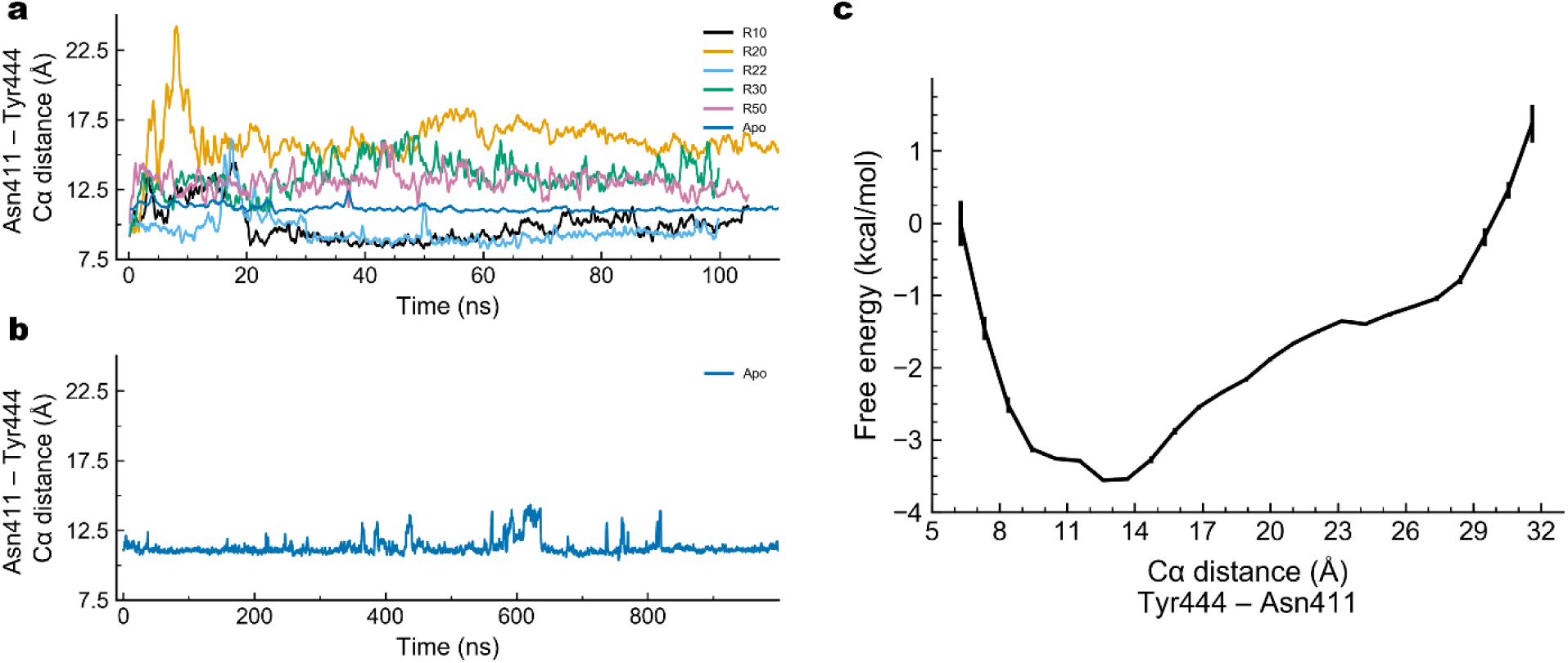
Lid-loop conformational changes. Cα distance between Asn411 and Tyr444 versus simulation time obtained **a**, from aMD runs and a conventional MD of apo EcoGGT. **b**, Extended conventional MD of apo EcoGGT. Data is smoothed by moving average over 20 frames – 400 ps for aMD runs and 800 ps for conventional MD. **c**, Free energy profile of the Tyr444 – Asn411 Cα-distance obtained from 60 aMD trajectories, ca. 6 microseconds in total. Error bars and data points represent standard deviation and averages calculated using 400 repeats of bootstrapping with replacement, respectively.

It is interesting to analyze the structure of the lid-loop when the substrate binding is rejected, namely trajectories R22 and R30, which showed near binding before repelling the substrate again into the solvent. R22 shows loop open conformations between ca. 15 ns to 25 ns, therefore blocking the substrate binding when it is near the pocket during the first 10 ns (Figures 2a-b, 4a-b, S8). In R30 the substrate fails to move into the pocket despite the lid-loop open conformation (Figure 4a). Our MD modeling indicates that the lid-loop acts as a steric barrier, enclosing the substrate in the pocket and points to a possible function as a gating element allowing the substrate to bind into the catalytic pocket. However, the lid opening is not the only condition necessary for the substrate binding. Favorable interactions between the pocket polar residues and the substrate that attracts and stabilize the substrate-enzyme complex is also essential as discussed in the previous subsection.

Phrases clustering showed that lid-loop residues form infrequent simultaneous contacts with the substrate. Instead, bound glutamine is frequently separated from the lid-loop by water molecules, as evident by calculating the number of water molecules between the substrate and Tyr444 (Figure S10). The number of water molecules are calculated using a moving box defined by the substrate and Tyr444 coordinates, further details and a VMD script are provided in Supplemental Material. Trajectories R20 and R50 have an average of 5.2 and 2.1 water molecules separating the substrate from the lid-loop over the last 15 ns, while R10 shows 0.4 water molecules. See final aMD snapshots in Figure S11.

### Structural Flexibility of the Glutamine-EcoGGT Binding Pocket

The conformational heterogeneity of the bound substrate and the catalytic site is analyzed by structural clustering over heavy-atom coordinates of the substrate and 14 residues located in the binding site, see Methods. This resulted in a set of conformations where the top 6 clusters amount for 58 % of the data set, and the top two clusters have a combined representation of 35 % of the data set (Table 1). A main difference between the two top clusters is an opening of the lid loop in cluster 2 measured by a 7 Å increase of the Cα distance of Tyr444 – Asn411 with respect to that distance (9 Å) in cluster 1, which results in a larger binding site entrance (Figures 5a-b). Note that, cluster 1 shows a conformational change where the N-terminus residue Thr391 is displaced by Pro482 (Figure 5a). Cluster 1 structures correspond to R10 trajectory. However, this conformational change is not observed in the second top cluster (encompassing structures from R20), in which the glutamine’s amide group is in close contact with Thr391. This shows that the different trajectories have heterogeneous substrate-bound complexes. Common interactions between the two top cluster representatives involve hydrogen bond interactions between Arg114 and Asn411 with the substrate and contacts with Gly484 (Figures 5c-d). Comparison between representative structures of clusters 3 and 4, both containing MD snapshots from R10, shows that residues in the binding pocket undergo subtle conformational changes that allow residues e.g. Ser462 and Ser463 to form hydrogen bonds with the substrate (Figure S12). Meanwhile, the lid-loop retains its conformation separating the binding pocket from the solvent (Figure S12). To further dissect the origins of the conformational flexibility, clustering analysis of substrate heavy-atom coordinates showed a dominant cluster accounting for ca. 80 % of the substrate-bound data set (Table S4). Thus, the substrate relative position and orientation and the contacting residues contribute the most to the substrate-pocket conformational heterogeneity, rather than the substrate conformational flexibility. Displacement of the lid-loop suggests that the loop acts as a gating entity by opening and closing the pocket entrance. Furthermore, conformational changes of pocket residues are possibly associated with the accommodation of different substrates into the catalytic reactive site. The implications of our modeling results will be further discussed in the Discussion and Conclusions section.

**Table 1.** Clustering analysis results of the conformational space of glutamine and binding pocket residues. The number of structures for each cluster is indicated. The cluster weight is calculated over the entire set of frames. The top six clusters represent 58.1 % of the data set. Details of clustering analysis are described in the Methods section.

| Cluster ID | Number of structures | Cluster population (%) |
| --- | --- | --- |
| 1 | 1842 | 21.7 |
| 2 | 1154 | 13.6 |
| 3 | 689 | 8.1 |
| 4 | 640 | 7.5 |
| 5 | 327 | 3.9 |
| 6 | 276 | 3.3 |

**Figure 5.**
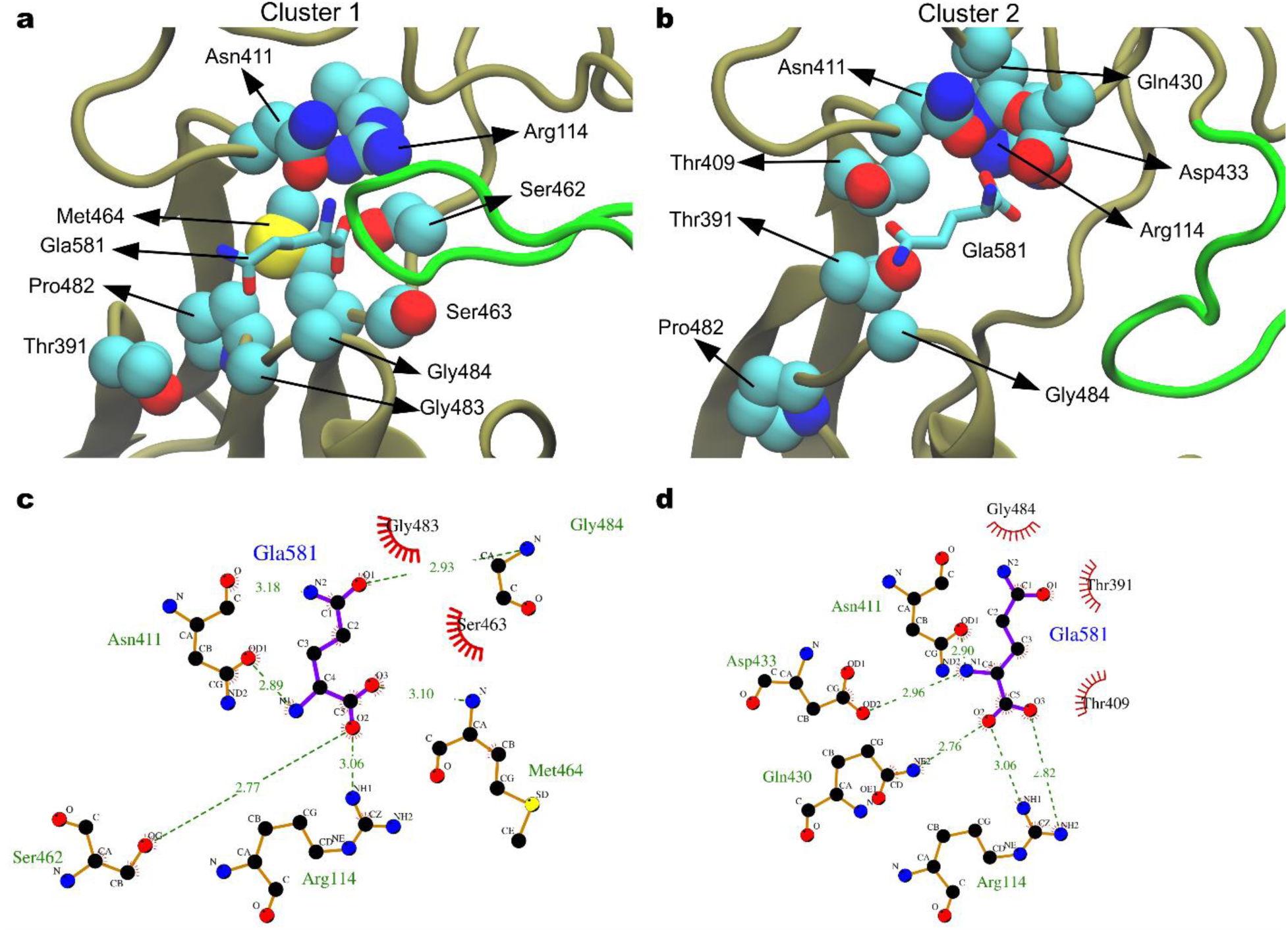
Top two cluster representative structures of the substrate (glutamine) bound to EcoGGT. **a**, Binding pocket of the representative structure of cluster 1. **b**, Hydrogen bond network and hydrophobic contacts of the glutamine (named as Gla581), and the protein pocket of cluster 1. **c**, and **d**, 2D scheme of binding pocket and substrate interactions of representative structure of cluster 1 and 2, respectively. The substrate is shown as ball and sticks and sidechains involved in hydrogen bonding and hydrophobic interactions are highlighted in licorice and space-filling representations. Lid loop (residues 439-448), is shown in green ribbon. Chain B is shown in tan ribbons. Nitrogen, carbon and oxygen atoms are represented in blue, cyan and red color spheres, respectively.

### Transient Complexes

We used conformational clustering on the substrate heavy-atom coordinates with the protein structure aligned relative to its initial conformation to identify transient complexes through the binding event observed in trajectory R20, selected as a representative aMD trajectory. Four cluster representatives out of five clusters are shown in Figure 6 and Supporting Movie S1. Cluster 2 is not shown since it is briefly populated after complete substrate binding (Figure S13). Clustering was carried out on the trajectory segment when the ligand RMSD is below 10 Å and performed with the GROMOS clustering method and cutoff radius of 3 Å (Figure S14). The first cluster (Figure 6a) attains a state common to all the five trajectories where glutamine approaches the binding pocket. The ligand is about 10 Å away from the catalytic residue and does not interact directly with the residues inside the active site. It is still able to depart from the protein surface, like in R22 and R30. This complex is also supported by the presence of residues that make hydrogen bonds with glutamine outside the binding pocket, namely Thr413 and Tyr194, as shown in the Specific Binding Site Interactions subsection. In the second cluster representative, glutamine is oriented in a way that allows it to make a stabilizing interaction between the negative charge on the carboxyl group and the N-terminus of Thr391. Glutamine’s amino group is also oriented towards the polar residue Asn411 and the negatively charged Asp433 (Figure 6b). The importance of these two residues was similarly highlighted in the previous subsections. In the third transient complex glutamine amino group interacts with Asn411 and Asp433. This interaction guides glutamine deep into the binding pocket (Figure 6c). Finally, in the last cluster the carboxyl group is engaged in hydrogen bonds with Arg114, Ser462 and Ser463 (residues in group I and II identified by the phrases clustering). The amino group is stabilized by Asn411 and Asp433. The amide group is now near the hydroxyl oxygen on Thr391 in a NAC conformation before the initial step of the reaction (Figure 6d).

**Figure 6.**
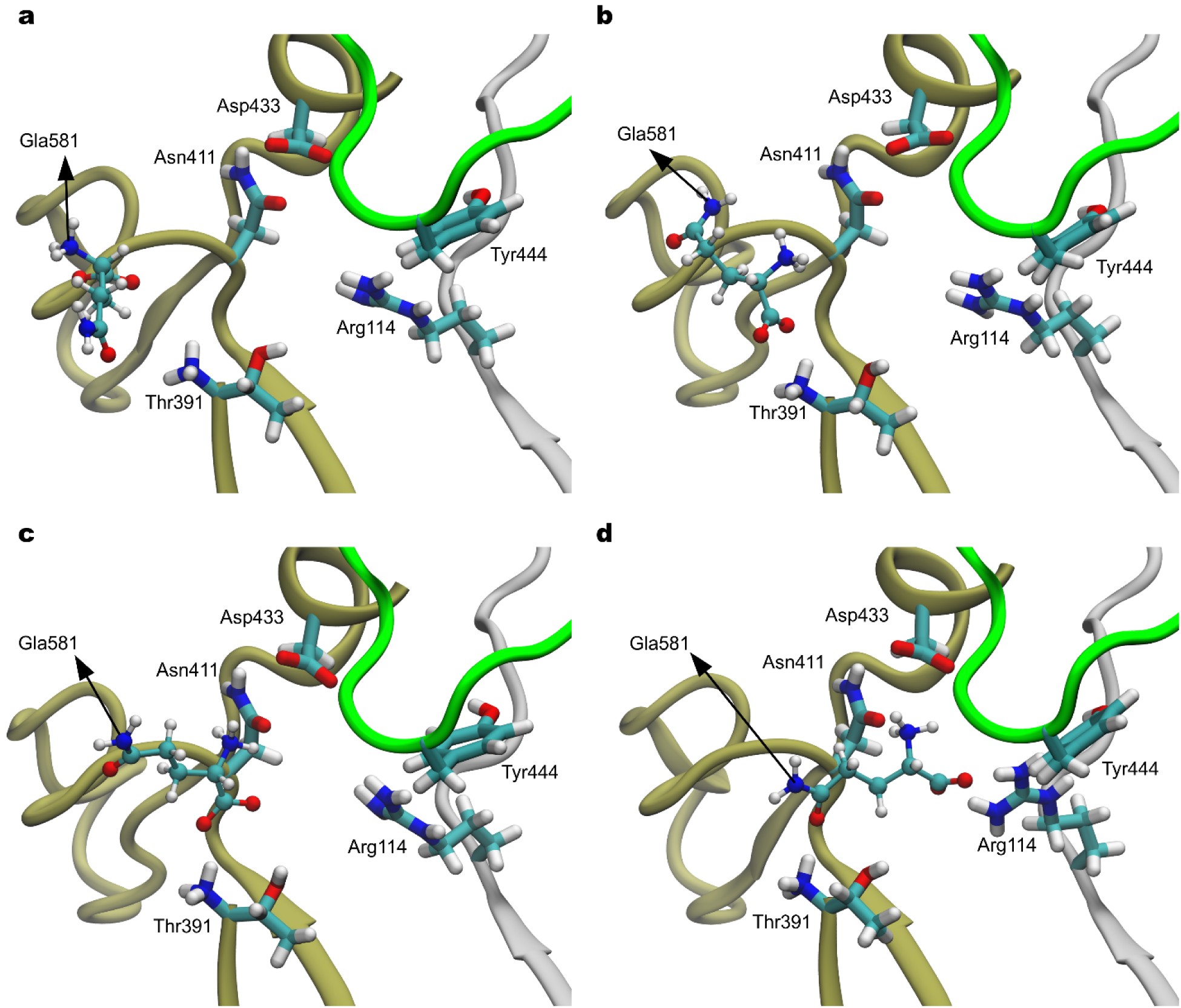
Cluster representative structures identified in the R20 trajectory segment. **a**, Cluster 4 **b**, cluster 5 **c**, cluster 3 and **d**, cluster 1. Glutamine (Gla581) is represented in licorice, the relevant residues in ball and sticks and the protein in ribbons representation. Nitrogen, carbon and oxygen atoms are represented in blue, cyan and red color spheres, respectively. White and magenta color ribbons represent the two subdomains of EcoGGT enzyme.

### Discussion and Conclusions

The process of substrate binding by enzymes may encompass key features of their activity and specificity. Here, a computational strategy based on a swarm of aMD simulations was employed to model the approach and subsequent binding of glutamine into the EcoGGT active site. Three binding events out of 60 independent short aMD repeats were observed, mirroring the actual behavior of enzymatic reactions, where only a small fraction of enzyme-substrate encounters results in effective binding events. Our simulated bound glutamine’s relative orientation and position is in good agreement with a naïvely built model of the EcoGGT bound to the substrate, based on a crystal X-ray structure of EcoGGT bound to the corresponding product obtained by Fukuyama et al.^24^ Based on MD postprocessing analysis, we identified transient complexes that could be relevant in the recognition process, imposing no restraint or knowledge based bias to the simulations.

Three structural motifs consisting of polar residues were found to stabilize through electrostatic interactions the substrate-pocket complex. One of the motifs contains residues Asp411 and Ser462 which form substrate contacts simultaneously with most other residues in the binding pocket. Based on our analysis, we hypothesize that electrostatic interactions between pocket residues and glutamine are the main driving force of substrate binding. This structural-dynamics insight could be employed for introducing polar/non-polar/aromatic substitutions in the identified binding motifs of EcoGGT, as further discussed below.

Our aMD simulations suggest that the lid-loop functions as a gating element and as a steric wall confining the substrate inside the binding pocket. The closed native state of the lid loop lies at the bottom of the free energy profile along the distance between the lid-loop tip and Asp411. Partially open states are accessible within 1 kT of the global free energy minimum (T = 300 K), while a fully open lid-loop state has a free energy cost of ca. 4 kT. This energetic description helps to interpret experimental insights, in which the lid loop’s role was probed through a variety of experimental techniques^59^. That work indicated that the loop acts through a gating mechanism allowing the selection of small polar substrates such as glutamine into the binding pocket of EcoGGT. This observation can be explained by the large free energy cost of a fully open loop conformation compared to partially open conformations accessible by thermal fluctuations. Overall, our simulations suggest that partially open conformations of the lid-loop are a necessary condition, but not the only one for the substrate to enter the binding site as favorable electrostatic interactions guide and stabilize the substrate in the binding pocket.

Another aspect revealed by our aMD simulations are conformational changes of residues in the pocket, which suggests pocket flexibility over longer timescales. Note that hundred-nanoseconds long aMD simulations were shown to reproduce protein fluctuations observed at millisecond timescale for several other proteins^60–62^. Furthermore, our cMD simulation of the apo state of EcoGGT showed a closed lid-loop conformation with short-lived displacements over 1 microsecond. This limited sampling obtained with conventional MD simulations justifies the use of swarm aMD simulations, an enhanced sampling method, to accelerate open/closed lid-loop transitions during the substrate binding and sample substrate-GGT bound structures.

Based in our molecular modelling, we propose a mixed model that explains key features of the GGT catalyzed reaction. Electrostatic interactions between the pocket amino acids and substrate govern the specificity of the enzymatic reaction and a conformational selection model that controls the binding through the relatively longer timescale of the enzyme conformational changes with respect to the protein-ligand diffusion^63,64^. A hypothesis is that the lid-loop gating mechanism is a general feature of GGT enzymes containing this conserved lid-loop and that specific site mutations on this loop may affect the enzyme selectivity. Targeting the flexibility of GGT’s lid-loop offers a strategy for the design of inhibitors that stabilize the loop^65^ or for the use of a mutational approach to enhance activity through single-, multiple-site, insertion and deletion of loop residues^66^. In conclusion, we proposed complexes of the glutamine-GGT molecular recognition process, and we also provided computational modeling evidence that supports a lid-loop gating mechanism governing substrate binding to EcoGGT. Even more, our all-atom simulations provide structural and dynamical insights for both the interpretation of GGT’s structure-function studies and the design of strategies targeting GGT’s activity and specificity features.

Finally, in this work we have introduced a computationally affordable protocol using the swarm aMD method to evaluate substrate binding, which could be used to study binding during the more complex GGT catalyzed transpeptidation reaction and potentially other ligand-protein systems.

## ASSOCIATED CONTENT

### Supporting Information

The following files are available free of charge. Supporting methodology details, tables and figures (PDF). Movie showing cluster representatives through the substrate binding pathway sampled by one aMD trajectory (MP4).

## AUTHOR INFORMATION

### Notes

The authors declare no competing financial interest.

## ACKNOWLEDGMENT

Computational calculations were performed at the Grendel-S cluster of the Centre for Scientific Computing Aarhus (CSCAA), Aarhus University and at the Abacus 2.0 of the DeIC National High-Performance Computing Centre, University of Southern Denmark.

## ABBREVIATIONS

GGT: γ-glutamyltransferase
EcoGGT: *Escherichia Coli* γ-glutamyltransferase
aMD: accelerated molecular dynamics

## REFERENCES

(1) Boehr, D. D.; Nussinov, R.; Wright, P. E. The Role of Dynamic Conformational Ensembles in Biomolecular Recognition. Nat. Chem. Biol. 2009, 5, 789–796.

(2) Lange, O. F.; Lakomek, N.-A.; Farès, C.; Schröder, G. F.; Walter, K. F. A.; Becker, S.; Meiler, J.; Grubmüller, H.; Griesinger, C.; Groot, B. L. de. Recognition Dynamics Up to Microseconds Revealed from an RDC-Derived Ubiquitin Ensemble in Solution. Science 2008, 320, 1471–1475.

(3) Eisenmesser, E. Z.; Millet, O.; Labeikovsky, W.; Korzhnev, D. M.; Wolf-Watz, M.; Bosco, D. A.; Skalicky, J. J.; Kay, L. E.; Kern, D. Intrinsic Dynamics of an Enzyme Underlies Catalysis. Nature 2005, 438, 117–121.

(4) Henzler-Wildman, K. A.; Lei, M.; Thai, V.; Kerns, S. J.; Karplus, M.; Kern, D. A Hierarchy of Timescales in Protein Dynamics Is Linked to Enzyme Catalysis. Nature 2007, 450, 913–916.

(5) Ahalawat, N.; Mondal, J. Mapping the Substrate Recognition Pathway in Cytochrome P450. J. Am. Chem. Soc. 2018, 140, 17743–17752.

(6) Shan, Y.; Kim, E. T.; Eastwood, M. P.; Dror, R. O.; Seeliger, M. A.; Shaw, D. E. How Does a Drug Molecule Find Its Target Binding Site? J. Am. Chem. Soc. 2011, 133, 9181–9183.

(7) Castellano, I.; Merlino, A. γ-Glutamyltranspeptidases: Sequence, Structure, Biochemical Properties, and Biotechnological Applications. Cell. Mol. Life Sci. 2012, 69, 3381–3394.

(8) Tate, S. S.; Meister, A. γ-Glutamyl Transpeptidase: Catalytic, Structural and Functional Aspects. Mol. Cell. Biochem. 1981, 39, 357–368.

(9) Ohkama-Ohtsu, N.; Oikawa, A.; Zhao, P.; Xiang, C.; Saito, K.; Oliver, D. J. A γ-Glutamyl Transpeptidase-Independent Pathway of Glutathione Catabolism to Glutamate via 5-Oxoproline in Arabidopsis. Plant Physiol. 2008, 148, 1603–1613.

(10) Ferretti, M.; Destro, T.; Tosatto, S. C. E.; La Rocca, N.; Rascio, N.; Masi, A. Gamma-Glutamyl Transferase in the Cell Wall Participates in Extracellular Glutathione Salvage from the Root Apoplast. New Phytol. 2009, 181, 115–126.

(11) Lieberman, M. W.; Wiseman, A. L.; Shi, Z. Z.; Carter, B. Z.; Barrios, R.; Ou, C. N.; Chévez-Barrios, P.; Wang, Y.; Habib, G. M.; Goodman, J. C.; Huang, S. L.; Lebovitz, R. M.; Matzuk, M. M. Growth Retardation and Cysteine Deficiency in Gamma-Glutamyl Transpeptidase-Deficient Mice. Proc. Natl. Acad. Sci. 1996, 93, 7923–7926.

(12) Hanigan, M. H.; Ricketts, W. A. Extracellular Glutathione Is a Source of Cysteine for Cells That Express .Gamma.-Glutamyl Transpeptidase. Biochemistry 1993, 32, 6302–6306.

(13) Mehdi, K.; Penninckx, M. J. An Important Role for Glutathione and γ-Glutamyltranspeptidase in the Supply of Growth Requirements during Nitrogen Starvation of the Yeast Saccharomyces Cerevisiae. Microbiology, 1997, 143, 1885–1889.

(14) Ricci, V.; Giannouli, M.; Romano, M.; Zarrilli, R. Helicobacter Pylori Gamma-Glutamyl Transpeptidase and Its Pathogenic Role. World J. Gastroenterol. 2014, 20, 630–638.

(15) Oinonen, C.; Rouvinen, J. Structural Comparison of Ntn-Hydrolases. Protein Sci. 2000, 9, 2329–2337.

(16) Inoue, M.; Hiratake, J.; Suzuki, H.; Kumagai, H.; Sakata, K. Identification of Catalytic Nucleophile of Escherichia Coli γ-Glutamyltranspeptidase by γ-Monofluorophosphono Derivative of Glutamic Acid: N-Terminal Thr-391 in Small Subunit Is the Nucleophile. Biochemistry 2000, 39, 7764–7771.

(17) Donald Allison, R. γ-Glutamyl Transpeptidase: Kinetics and Mechanism. Methods Enzymol. 1985, 113, 419–437.

(18) Taniguchi, N.; Ikeda, Y. γ-Glutamyl Transpeptidase: Catalytic Mechanism and Gene Expression. In Advances in Enzymology and Related Areas of Molecular Biology; John Wiley & Sons, Ltd, 2006; pp 239–278.

(19) Tate, S. S.; Meister, A. Interaction of γ-Glutamyl Transpeptidase with Amino Acids, Dipeptides, and Derivatives and Analogs of Glutathione. J. Biol. Chem. 1974, 249, 7593–7602.

(20) Suzuki, H.; Hashimoto, W.; Kumagai, H. Escherichia Coli K-12 Can Utilize an Exogenous Gamma-Glutamyl Peptide as an Amino Acid Source, for Which Gamma-Glutamyltranspeptidase Is Essential. J. Bacteriol. 1993, 175, 6038–6040.

(21) Suzuki, H.; Yamada, C.; Kato, K. γ-Glutamyl Compounds and Their Enzymatic Production Using Bacterial γ-Glutamyltranspeptidase. Amino Acids 2007, 32, 333–340.

(22) Suzuki, H.; Kumagai, H.; Tochikura, T. Gamma-Glutamyltranspeptidase from Escherichia Coli K-12: Purification and Properties. J. Bacteriol. 1986, 168, 1325–1331.

(23) Sakai, H.; Sakabe, N.; Sasaki, K.; Hashimoto, W.; Suzuki, H.; Tachi, H.; Kumagai, H.; Sakabe, K. A Preliminary Description of the Crystal Structure of γ-Glutamyltranspeptidase from E. Coli K-12. J. Biochem. (Tokyo) 1996, 120, 26–28.

(24) Kumagai, H.; Suzuki, H.; Okada, T.; Wada, K.; Fukuyama, K. Crystal Structures of γ-Glutamyltranspeptidase from Escherichia Coli, a Key Enzyme in Glutathione Metabolism, and Its Reaction Intermediate. Proc. Natl. Acad. Sci. 2006, 103, 6471–6476.

(25) Suzuki, H.; Kumagai, H. Autocatalytic Processing of γ-Glutamyltranspeptidase. J. Biol. Chem. 2002, 277, 43536–43543.

(26) Hiratake, J.; Okada, T.; Kumagai, H.; Wada, K.; Yamada, C.; Irie, M.; Suzuki, H.; Fukuyama, K. Crystal Structures of Escherichia Coli γ-Glutamyltranspeptidase in Complex with Azaserine and Acivicin: Novel Mechanistic Implication for Inhibition by Glutamine Antagonists. J. Mol. Biol. 2008, 380, 361–372.

(27) West, M. B.; Chen, Y.; Wickham, S.; Heroux, A.; Cahill, K.; Hanigan, M. H.; Mooers, B. H. M. Novel Insights into Eukaryotic γ-Glutamyltranspeptidase 1 from the Crystal Structure of the Glutamate-Bound Human Enzyme. J. Biol. Chem. 2013, 288, 31902–31913.

(28) Pan, A. C.; Borhani, D. W.; Dror, R. O.; Shaw, D. E. Molecular Determinants of Drug– Receptor Binding Kinetics. Drug Discov. Today 2013, 18, 667–673.

(29) Hollingsworth, S. A.; Dror, R. O. Molecular Dynamics Simulation for All. Neuron 2018, 99, 1129–1143.

(30) Best, R. B. Atomistic Molecular Simulations of Protein Folding. Curr. Opin. Struct. Biol. 2012, 22, 52–61.

(31) Sugita, Y.; Okamoto, Y. Replica-Exchange Molecular Dynamics Method for Protein Folding. Chem. Phys. Lett. 1999, 314, 141–151.

(32) Laio, A.; Parrinello, M. Escaping Free-Energy Minima. Proc. Natl. Acad. Sci. U. S. A. 2002, 99, 12562–12566.

(33) Torrie, G. M.; Valleau, J. P. Monte Carlo Free Energy Estimates Using Non-Boltzmann Sampling: Application to the Sub-Critical Lennard-Jones Fluid. Chem. Phys. Lett. 1974, 28, 578–581.

(34) Dellago, C.; Bolhuis, P. G.; Csajka, F. S.; Chandler, D. Transition Path Sampling and the Calculation of Rate Constants. J. Chem. Phys. 1998, 108, 1964–1977.

(35) Hamelberg, D.; Mongan, J.; McCammon, J. A. Accelerated Molecular Dynamics: A Promising and Efficient Simulation Method for Biomolecules. J. Chem. Phys. 2004, 120, 11919–11929.

(36) Doshi, U.; Hamelberg, D. Achieving Rigorous Accelerated Conformational Sampling in Explicit Solvent. J. Phys. Chem. Lett. 2014, 5, 1217–1224.

(37) Flores-Canales, J. C.; Kurnikova, M. Targeting Electrostatic Interactions in Accelerated Molecular Dynamics with Application to Protein Partial Unfolding. J. Chem. Theory Comput. 2015, 11, 2550–2559.

(38) Miao, Y.; Feher, V. A.; McCammon, J. A. Gaussian Accelerated Molecular Dynamics: Unconstrained Enhanced Sampling and Free Energy Calculation. J. Chem. Theory Comput. 2015, 11, 3584–3595.

(39) Miao, Y.; McCammon, J. A. Graded Activation and Free Energy Landscapes of a Muscarinic G-Protein–Coupled Receptor. Proc. Natl. Acad. Sci. 2016, 113, 12162–12167.

(40) Kappel, K.; Miao, Y.; McCammon, J. A. Accelerated Molecular Dynamics Simulations of Ligand Binding to a Muscarinic G-Protein-Coupled Receptor. Q. Rev. Biophys. 2015, 48, 479–487.

(41) Andersen, O. J.; Risør, M. W.; Poulsen, E. C.; Nielsen, N. Chr.; Miao, Y.; Enghild, J. J.; Schiøtt, B. Reactive Center Loop Insertion in α-1-Antitrypsin Captured by Accelerated Molecular Dynamics Simulation. Biochemistry 2017, 56, 634–646.

(42) Hamelberg, D.; de Oliveira, C. A. F.; McCammon, J. A. Sampling of Slow Diffusive Conformational Transitions with Accelerated Molecular Dynamics. J. Chem. Phys. 2007, 127, 155102.

(43) Hornak, V.; Okur, A.; Rizzo, R. C.; Simmerling, C. HIV-1 Protease Flaps Spontaneously Open and Reclose in Molecular Dynamics Simulations. Proc. Natl. Acad. Sci. 2006, 103, 915–920.

(44) Jorgensen, W. L.; Chandrasekhar, J.; Madura, J. D.; Impey, R. W.; Klein, M. L. Comparison of Simple Potential Functions for Simulating Liquid Water. J. Chem. Phys. 1983, 79, 926–935.

(45) Eastman, P.; Swails, J.; Chodera, J. D.; McGibbon, R. T.; Zhao, Y.; Beauchamp, K. A.; Wang, L.-P.; Simmonett, A. C.; Harrigan, M. P.; Stern, C. D.; Wiewiora, R. P.; Brooks, B. R.; Pande, V. S. OpenMM 7: Rapid Development of High Performance Algorithms for Molecular Dynamics. PLOS Comput. Biol. 2017, 13, e1005659.

(46) Miyamoto, S.; Kollman, P. A. Settle: An Analytical Version of the SHAKE and RATTLE Algorithm for Rigid Water Models. J. Comput. Chem. 1992, 13, 952–962.

(47) Vanden-Eijnden, E.; Ciccotti, G. Second-Order Integrators for Langevin Equations with Holonomic Constraints. Chem. Phys. Lett. 2006, 429, 310–316.

(48) Chow, K.-H.; Ferguson, D. M. Isothermal-Isobaric Molecular Dynamics Simulations with Monte Carlo Volume Sampling. Comput. Phys. Commun. 1995, 91, 283–289.

(49) Åqvist, J.; Wennerström, P.; Nervall, M.; Bjelic, S.; Brandsdal, B. O. Molecular Dynamics Simulations of Water and Biomolecules with a Monte Carlo Constant Pressure Algorithm. Chem. Phys. Lett. 2004, 384, 288–294.

(50) Lindert, S.; Bucher, D.; Eastman, P.; Pande, V.; McCammon, J. A. Accelerated Molecular Dynamics Simulations with the AMOEBA Polarizable Force Field on Graphics Processing Units. J. Chem. Theory Comput. 2013, 9, 4684–4691.

(51) Kjølbye, L. R.; Laustsen, A.; Vestergaard, M.; Periole, X.; De Maria, L.; Svendsen, A.; Coletta, A.; Schiøtt, B. Molecular Modeling Investigation of the Interaction between *Humicola Insolens* Cutinase and SDS Surfactant Suggests a Mechanism for Enzyme Inactivation. J. Chem. Inf. Model. 2019, 59, 1977–1987.

(52) Arnarez, C.; Mazat, J.-P.; Elezgaray, J.; Marrink, S.-J.; Periole, X. Evidence for Cardiolipin Binding Sites on the Membrane-Exposed Surface of the Cytochrome Bc1. J. Am. Chem. Soc. 2013, 135, 3112–3120.

(53) Abraham, M. J.; Murtola, T.; Schulz, R.; Páll, S.; Smith, J. C.; Hess, B.; Lindahl, E. GROMACS: High Performance Molecular Simulations through Multi-Level Parallelism from Laptops to Supercomputers. SoftwareX 2015, 1–2, 19–25.

(54) Daura, X.; Gademann, K.; Jaun, B.; Seebach, D.; van Gunsteren, W. F.; Mark, A. E. Peptide Folding: When Simulation Meets Experiment. Angew. Chem. Int. Ed. 1999, 38, 236–240.

(55) Humphrey, W.; Dalke, A.; Schulten, K. VMD: Visual Molecular Dynamics. J. Mol. Graph. 1996, 14, 33–38.

(56) Laskowski, R. A.; Swindells, M. B. LigPlot+: Multiple Ligand–Protein Interaction Diagrams for Drug Discovery. J. Chem. Inf. Model. 2011, 51, 2778–2786.

(57) Miao, Y.; Sinko, W.; Pierce, L.; Bucher, D.; Walker, R. C.; McCammon, J. A. Improved Reweighting of Accelerated Molecular Dynamics Simulations for Free Energy Calculation. J. Chem. Theory Comput. 2014, 10, 2677–2689.

(58) Bruice, T. C.; Lightstone, F. C. Ground State and Transition State Contributions to the Rates of Intramolecular and Enzymatic Reactions. Acc. Chem. Res. 1999, 32, 127–136.

(59) Calvio, C.; Romagnuolo, F.; Vulcano, F.; Speranza, G.; Morelli, C. F. Evidences on the Role of the Lid Loop of γ-Glutamyltransferases (GGT) in Substrate Selection. Enzyme Microb. Technol. 2018, 114, 55–62.

(60) Pierce, L. C. T.; Salomon-Ferrer, R.; Augusto F. de Oliveira, C.; McCammon, J. A.; Walker, R. C. Routine Access to Millisecond Time Scale Events with Accelerated Molecular Dynamics. J. Chem. Theory Comput. 2012, 8, 2997–3002.

(61) Markwick, P. R. L.; Bouvignies, G.; Blackledge, M. Exploring Multiple Timescale Motions in Protein GB3 Using Accelerated Molecular Dynamics and NMR Spectroscopy. J. Am. Chem. Soc. 2007, 129, 4724–4730.

(62) Miao, Y.; Nichols, S. E.; Gasper, P. M.; Metzger, V. T.; McCammon, J. A. Activation and Dynamic Network of the M2 Muscarinic Receptor. Proc. Natl. Acad. Sci. 2013, 110, 10982–10987.

(63) McCammon, J. A.; Northrup, S. H. Gated Binding of Ligands to Proteins. Nature 1981, 293, 316–317.

(64) Zhou, H.-X. From Induced Fit to Conformational Selection: A Continuum of Binding Mechanism Controlled by the Timescale of Conformational Transitions. Biophys. J. 2010, 98, L15–L17.

(65) Suebsuwong, C.; Pinkas, D. M.; Ray, S. S.; Bufton, J. C.; Dai, B.; Bullock, A. N.; Degterev, A.; Cuny, G. D. Activation Loop Targeting Strategy for Design of Receptor-Interacting Protein Kinase 2 (RIPK2) Inhibitors. Bioorg. Med. Chem. Lett. 2018, 28, 577–583.

(66) Nestl, B. M.; Hauer, B. Engineering of Flexible Loops in Enzymes. ACS Catal. 2014, 4, 3201–3211.

