## Supplementary material for "Simulating Multiple Substrate Binding Events by γ-Glutamyltransferase using Accelerated Molecular Dynamics": supp_info

**Accelerated Molecular Dynamics (aMD):**

The potential in a aMD^1^ simulation is calculated as follows

$$V^{*}\left( r \right)=\left\{ \begin{aligned} V\left( r \right)+\Delta V,V\left( r \right)<E \\ V\left( r \right),V\left( r \right)\geq E \end{aligned} \right.$$

Where $V^{*}\left( r \right)$ is the modified potential, V(r) is the real potential and E is the energy threshold.

And $\Delta V$

$$\Delta V\left( r \right)=\left\{ \begin{aligned} \frac{\left( E-V(r) \right)^{2}}{\alpha+(E-V(r)}, &V(r)<0 \\ 0, &V(r)\geq0 \end{aligned} \right.$$

Where $\alpha$ is the acceleration parameter.

In the dual boost^2^ approach, the real potential is split in two different parameter

$$V\left( r \right)= V_{0}\left( r \right)+V_{t}(r)$$

Where $V_{t}\left( r \right)$ is the total contribution term and $V_{0}\left( r \right)$ is the sum of all the others contributions.

The first boost is the boost to the torsional term, ${\Delta V}_{t}(r)$. The resulting modified potential is called total potential, $V_{T}\left( r \right)$.

$$V_{T}\left( r \right)=V_{0}\left( r \right)+V_{t}\left( r \right)+{\Delta V}_{t}(r)$$

A secondary boost is then added to the total potential

$$V^{*}\left( r \right)=V_{0}\left( r \right)+V_{t}\left( r \right)+{\Delta V}_{t}\left( r \right)+{\Delta V}_{T}\left( r \right)$$

Is now possible to calculate the force on the modified potential

$$F^{*}=-\nabla V^{*}\left( r \right)$$

Therefore

$$F^{*}=-\left\{ \nabla V_{0}\left( r \right)+\nabla V_{t}\left( r \right)\left( \frac{\alpha_{t}}{\alpha_{t}+E_{t}-V_{t}(r)} \right)^{2} \right\}\times\left( \frac{\alpha_{T}}{\alpha_{T}+E_{T}-V_{T}(r)} \right)^{2}$$

The parameters, $\alpha_{t}$, $E_{t}$, $\alpha_{T}$, $E_{T}$, are the torsional boost energy and the total boost energy, respectively and are be calculated as follows.

$E_{t}= V_{t}+ a_{1} N_{res}$	$\alpha_{t}=a_{2}\frac{N_{res}}{5}$

$E_{T}= V_{T}+ b_{1} N_{atoms}$	$\alpha_{T}=b_{2}N_{atoms}$

$a_{1}$, $a_{2}$, $b_{1}$, $b_{2}$, are coefficient reported by Miao, et al^3^ and are $a_{1}$ = $a_{2}$ = 3.5 and $b_{1}$= $b_{2}$ = 0.175. $E_{t}$ and $E_{T}$ are calculated after a short cMD (50 ns) and were set $E_{t}=5662.3$ kCal·mol and $E_{T}=-273982.6$ kCal·mol

**Contact analysis:**

To further investigate how glutamine approaches the binding pocket, we analyzed the contacts established between glutamine and protein residues in the successful trajectories (R10, R20 and R50), identifying which ones were common to all of them. Since the time spent by glutamine in the binding pocket is different for each trajectory, we normalized the persistence time of each contact with respect to the highest one for each MD run.

It is interesting to notice that the residue that always scores the highest amount of contacts is Asn 411, which is used to normalize the number of contacts. Normalized contact percentages are defined as $Normalized Contacts= C_{R}\cdot100/C_{N411}$, where $C_{R}$ is the number of contacts between glutamine and a specific residue. $C_{N411}$ is the number of contacts between glutamine and Asn 411. Contacts are considered between a pair of heavy-atoms with a cutoff distance of 3.0 Å. We considered in further analyses only contacts that were common to all the three successful trajectories and were observed for at least 5% of the normalized contacts population (Tables S1-S3). Particularly important residues are Arg114, Thr409 and Asp433, which interact with glutamine for more than 30% of the normalized contacts in each of the three trajectories. This is also shown in plots of contact formation as function of time plots (Figs. S1-S6). We calculated the contacts between glutamine and EcoGGT using a plugin freely available and authored by Lubos Vrbka. The results where then analyzed using a python script.

| R10 | | | R20 | | | R50 | | | |
| --- | --- | --- | --- | --- | --- | --- | --- | --- | --- |
| Residue | Number of contacts | Percentage (%) | Residue | Number of contacts | Percentage (%) | Residue | Number of contacts | Percentage (%) | |
| 114ARG | 3472 | 74.67 | 114ARG | 1839 | 100.00 | 114ARG | 841 | | 34.95 |
| 192LYS | 5 | 0.11 | 192LYS | 2 | 0.11 | 192LYS | 17 | | 0.71 |
| 194TYR | 28 | 0.60 | 194TYR | 20 | 1.09 | 194TYR | 76 | | 3.16 |
| 218LYS | 5 | 0.11 | 218LYS | 13 | 0.71 | 218LYS | 11 | | 0.46 |
| 391THR | 323 | 6.95 | 391THR | 1758 | 95.60 | 391THR | 1548 | | 64.34 |
| 409THR | 3455 | 74.30 | 409THR | 1762 | 95.81 | 409THR | 1861 | | 77.35 |
| 411ASN | 4650 | 100.00 | 411ASN | 1839 | 100.00 | 411ASN | 2406 | | 100.00 |
| 412THR | 371 | 7.98 | 412THR | 45 | 2.45 | 412THR | 473 | | 19.66 |
| 413THR | 73 | 1.57 | 413THR | 17 | 0.92 | 413THR | 146 | | 6.07 |
| 430GLN | 7 | 0.15 | 430GLN | 1676 | 91.14 | 430GLN | 876 | | 36.41 |
| 433ASP | 1851 | 39.81 | 433ASP | 1571 | 85.43 | 433ASP | 1727 | | 71.78 |
| 444TYR | 1308 | 28.13 | 444TYR | 27 | 1.47 | 444TYR | 1755 | | 72.94 |
| 461LEU | 42 | 0.90 | 461LEU | 2 | 0.11 | 461LEU | 6 | | 0.25 |
| 462SER | 4289 | 92.24 | 462SER | 1017 | 55.30 | 462SER | 14 | | 0.58 |
| 463SER | 4401 | 94.65 | 463SER | 133 | 7.23 | 463SER | 135 | | 5.61 |
| 464MET | 3753 | 80.71 | 464MET | 369 | 20.07 | 464MET | 188 | | 7.81 |
| 483GLY | 2892 | 62.19 | 483GLY | 15 | 0.82 | 483GLY | 96 | | 3.99 |
| 484GLY | 4140 | 89.03 | 484GLY | 1385 | 75.31 | 484GLY | 1189 | | 49.42 |
| 485SER | 154 | 3.31 | 485SER | 113 | 6.14 | 485SER | 975 | | 40.52 |
| 487ILE | 2 | 0.04 | 487ILE | 48 | 2.61 | 487ILE | 47 | | 1.95 |
| 547LYS | 62 | 1.33 | 547LYS | 15 | 0.82 | 547LYS | 22 | | 0.91 |
| 573VAL | 16 | 0.34 | 573VAL | 17 | 0.92 | 573VAL | 2 | | 0.08 |

**Table S1.** Contact analysis results for aMD trajectories where substrate binds stably to the receptor. The table shows the absolute number of contacts and the percentage averaged over 411ASN’s number of contacts. Only the common residues are shown

| R10 | | | R20 | | | R50 | | |
| --- | --- | --- | --- | --- | --- | --- | --- | --- |
| Residue | Number of contacts | Percentage (%) | Residue | Number of contacts | Percentage (%) | Residue | Number of contacts | Percentage (%) |
| 114ARG | 3472 | 74.67 | 114ARG | 1839 | 100.00 | 114ARG | 841 | 34.95 |
| 391THR | 323 | 6.95 | 391THR | 1758 | 95.60 | 391THR | 1548 | 64.34 |
| 409THR | 3455 | 74.30 | 409THR | 1762 | 95.81 | 409THR | 1861 | 77.35 |
| 411ASN | 4650 | 100.00 | 411ASN | 1839 | 100.00 | 411ASN | 2406 | 100.00 |
| 412THR | 371 | 7.98 | 412THR | 45 | 2.45 | 412THR | 473 | 19.66 |
| 413THR | 73 | 1.57 | 413THR | 17 | 0.92 | 413THR | 146 | 6.07 |
| 430GLN | 7 | 0.15 | 430GLN | 1676 | 91.14 | 430GLN | 876 | 36.41 |
| 433ASP | 1851 | 39.81 | 433ASP | 1571 | 85.43 | 433ASP | 1727 | 71.78 |
| 444TYR | 1308 | 28.13 | 444TYR | 27 | 1.47 | 444TYR | 1755 | 72.94 |
| 462SER | 4289 | 92.24 | 462SER | 1017 | 55.30 | 462SER | 14 | 0.58 |
| 463SER | 4401 | 94.65 | 463SER | 133 | 7.23 | 463SER | 135 | 5.61 |
| 464MET | 3753 | 80.71 | 464MET | 369 | 20.07 | 464MET | 188 | 7.81 |
| 483GLY | 2892 | 62.19 | 483GLY | 15 | 0.82 | 483GLY | 96 | 3.99 |
| 484GLY | 4140 | 89.03 | 484GLY | 1385 | 75.31 | 484GLY | 1189 | 49.42 |
| 485SER | 154 | 3.31 | 485SER | 113 | 6.14 | 485SER | 975 | 40.52 |

**Table S2.** Contact analysis results for aMD trajectories where substrate binds stably to the receptor. Same data showed in Table S1 but only the residues that scores more than 5% in at least one of the three trajectories are shown.

| Residue | Average % | |
| --- | --- | --- |
| Arg114 | | 69.87 |
| Thr391 | | 55.63 |
| Thr409 | | 82.49 |
| Asn411 | | 100.00 |
| Thr412 | | 10.03 |
| Thr413 | | 2.85 |
| Gln430 | | 42.57 |
| Asp433 | | 65.67 |
| Tyr444 | | 34.18 |
| Ser462 | | 49.37 |
| Ser463 | | 35.83 |
| Met464 | | 36.20 |
| Gly483 | | 22.33 |
| Gly484 | | 71.25 |
| Ser485 | | 16.66 |

**Table S3.** Normalized Contacts average of three successful trajectories R10, R20 and R50. The averages are calculated as sum of the normalized percentages divided by three.

| Cluster | Number of structures | Cluster weight (%) | RMSD (Å) |
| --- | --- | --- | --- |
| 1 | 6775 | 79.9 | 0.91 |
| 2 | 1045 | 12.3 | 1.50 |
| 3 | 309 | 3.6 | 1.05 |
| 4 | 106 | 1.2 | 2.96 |

**Table S4.** Clustering of the conformational space sampled by glutamine after binding EcoGGT. The number of structures for each cluster is indicated. The cluster weight is calculated over the entire set of clusters. The RMSD is calculated relative to the proposed complex protein-glutamine structure after converting the glutamate into glutamine. Clustering analysis details are provided in the Methods section.

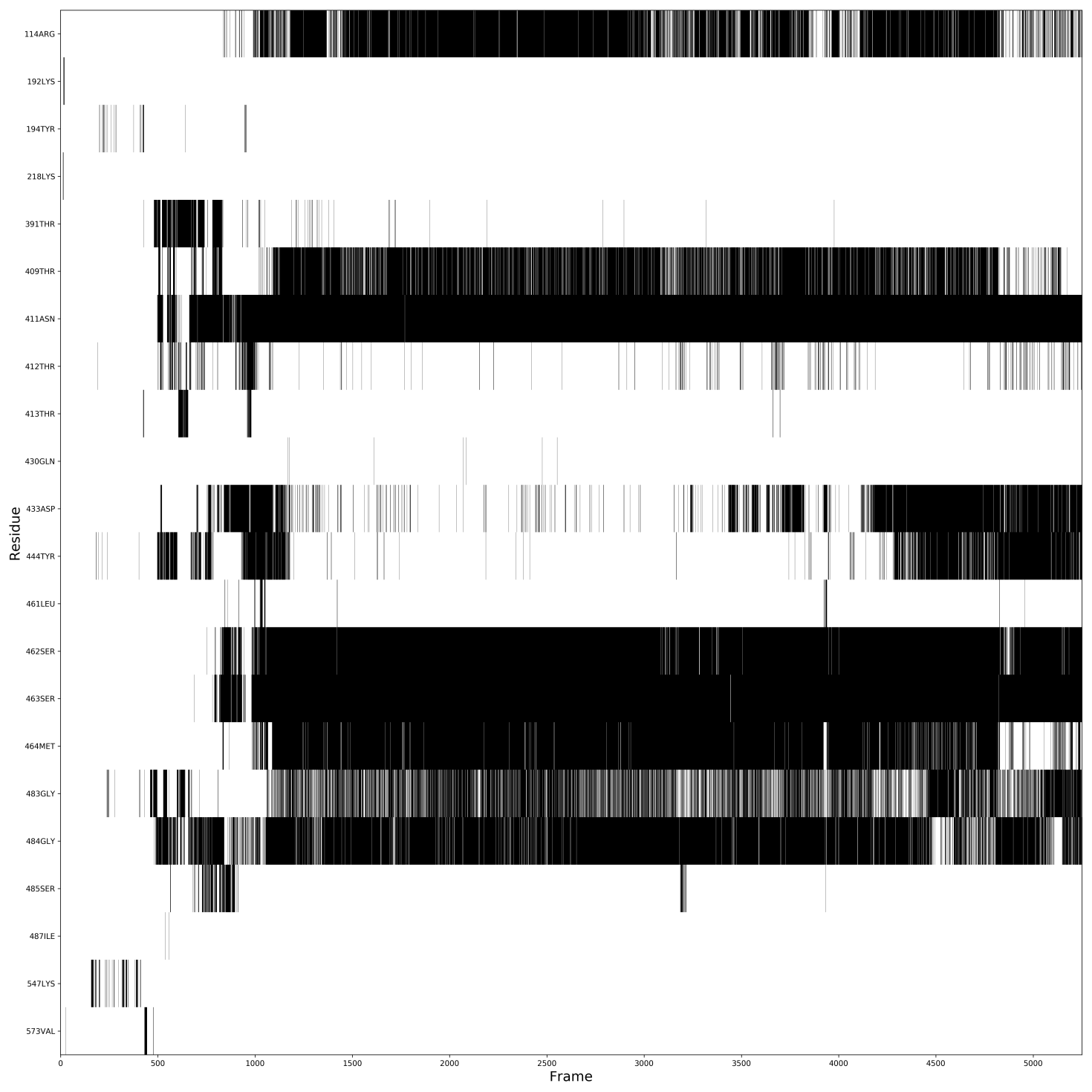
**Figure S1.** Contact analysis over time in aMD trajectory R10. Only the common contacts are shown.
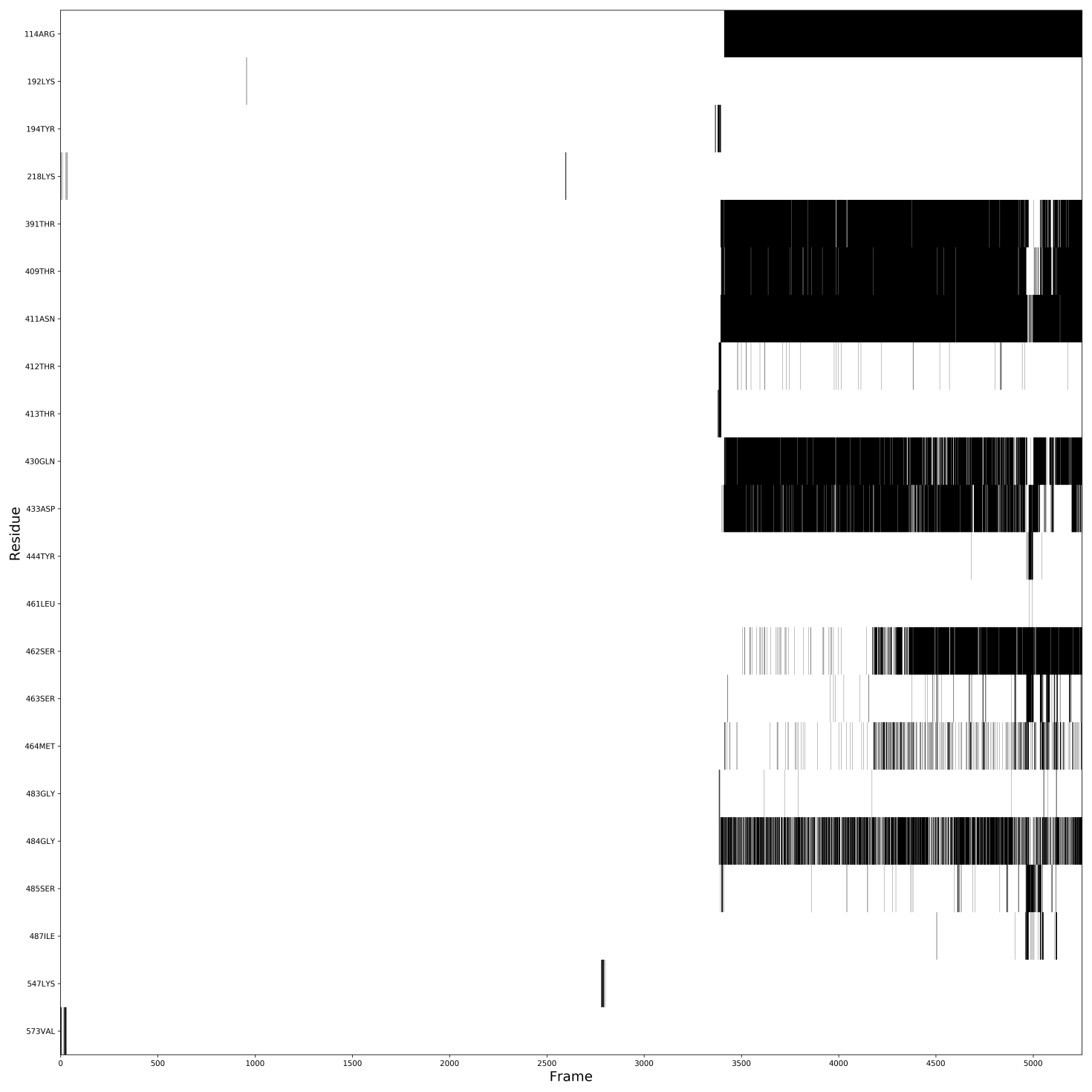
 **Figure S2.** Contact analysis over time in aMD trajectory R20. Only the common contacts are shown.
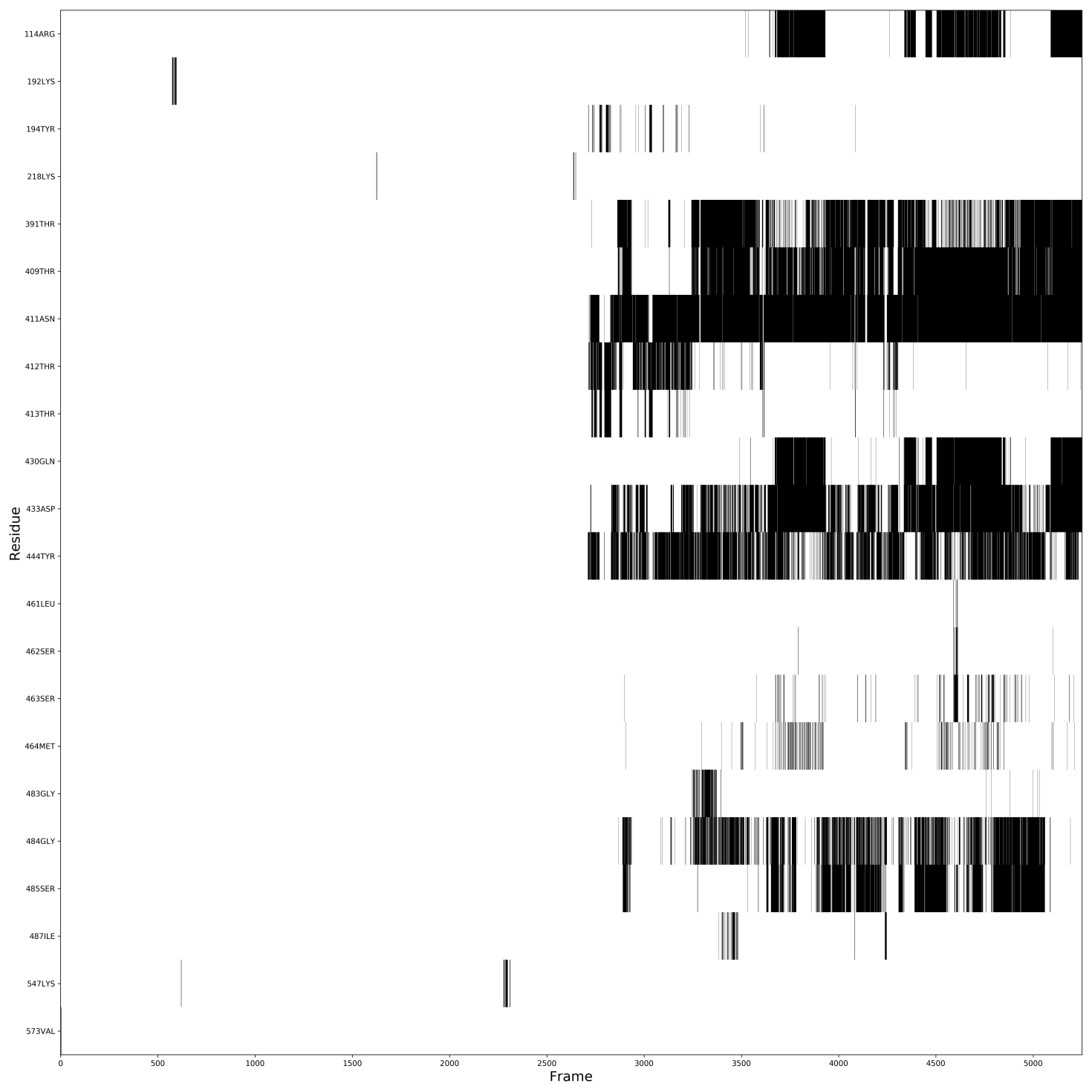
 **Figure S3.** Contact analysis over time in aMD trajectory R50. Only the common contacts are shown.
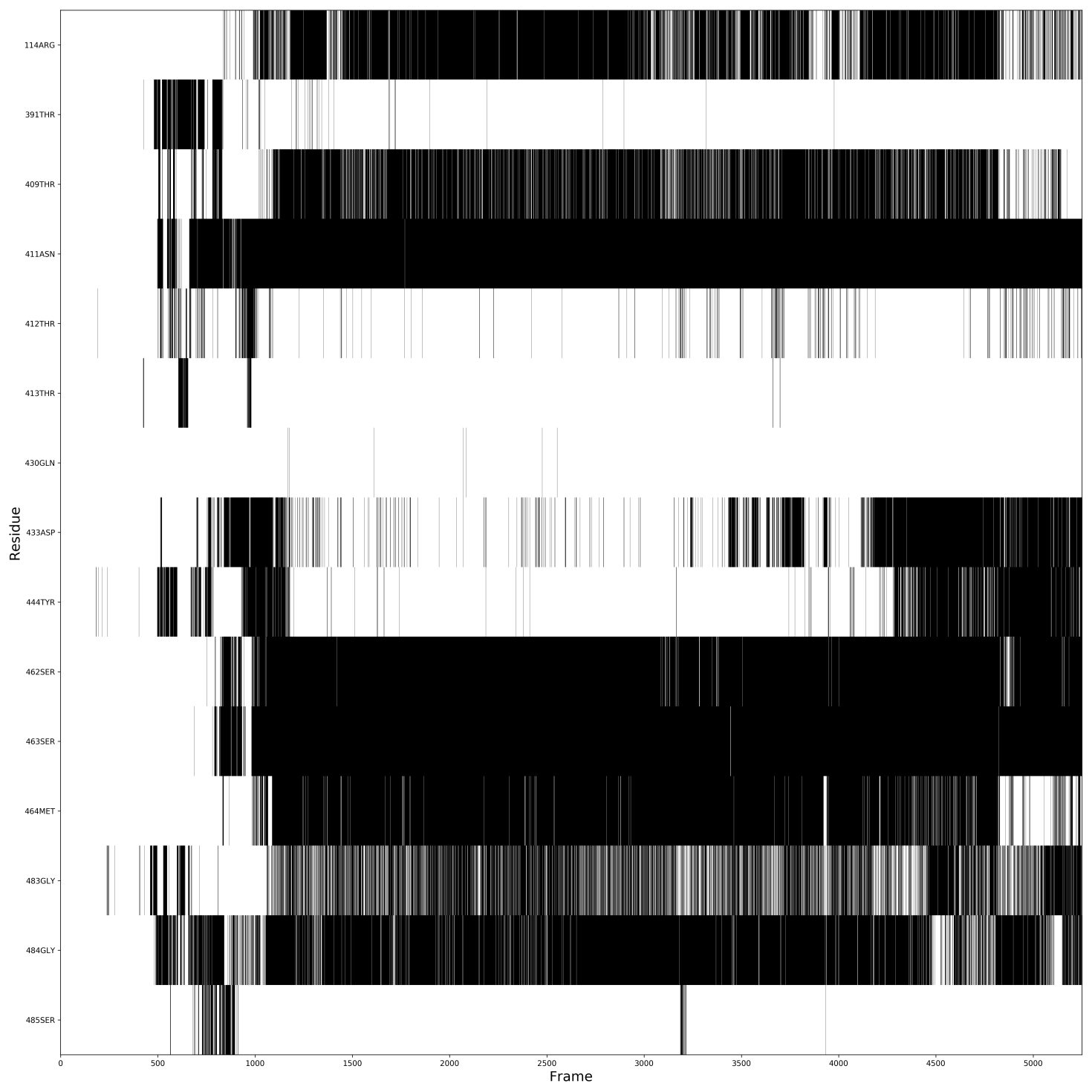
 **Figure S4.** Contact analysis over time in aMD trajectory R10. Only the residues calculated in Table S2 are shown.
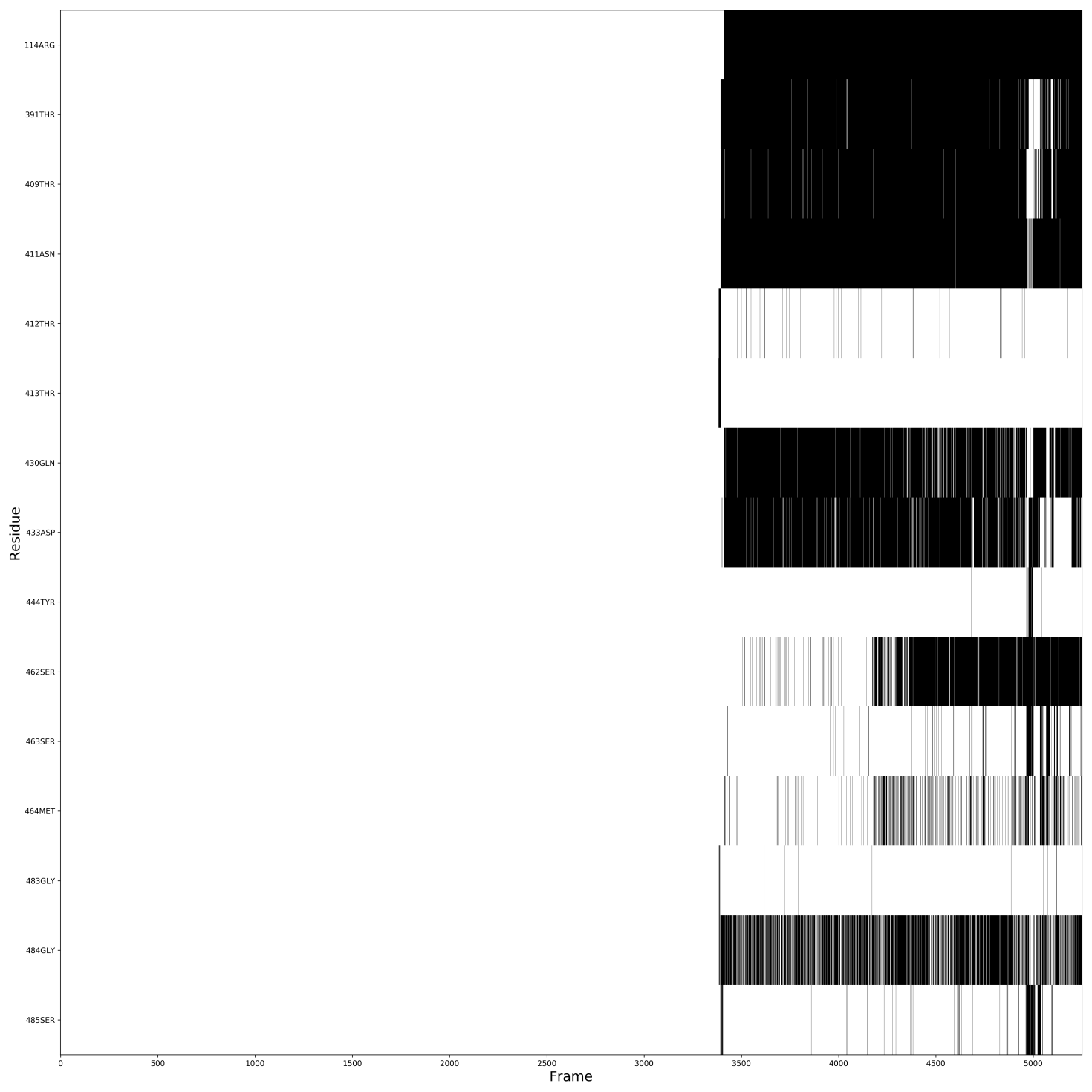
 **Figure S5.** Contact analysis over time in aMD trajectory R20. Only the residues calculated in Table S2 are shown.
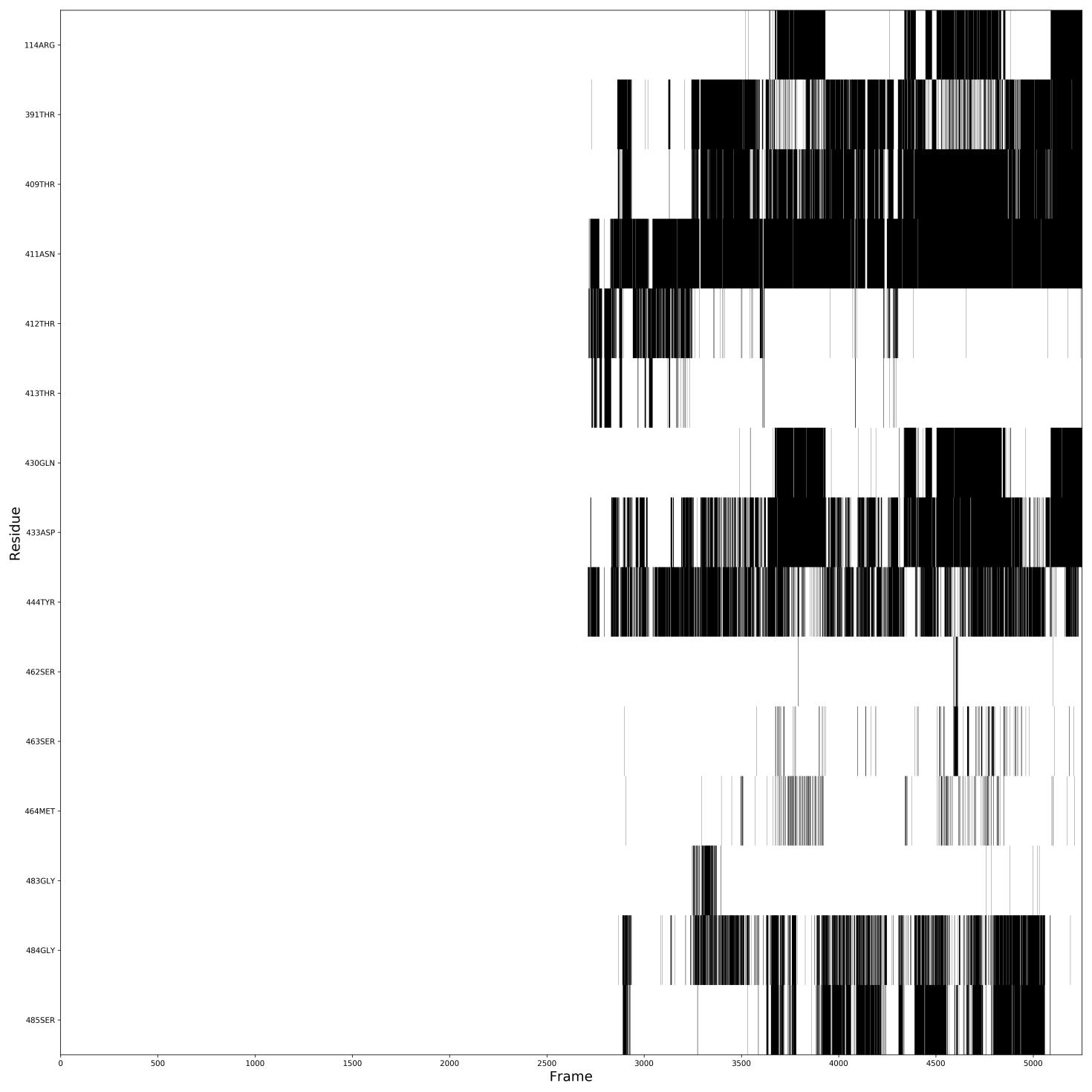
 **Figure S6.** Contact analysis over time in aMD trajectory R50. Only the residues calculated in Table S2 are shown.

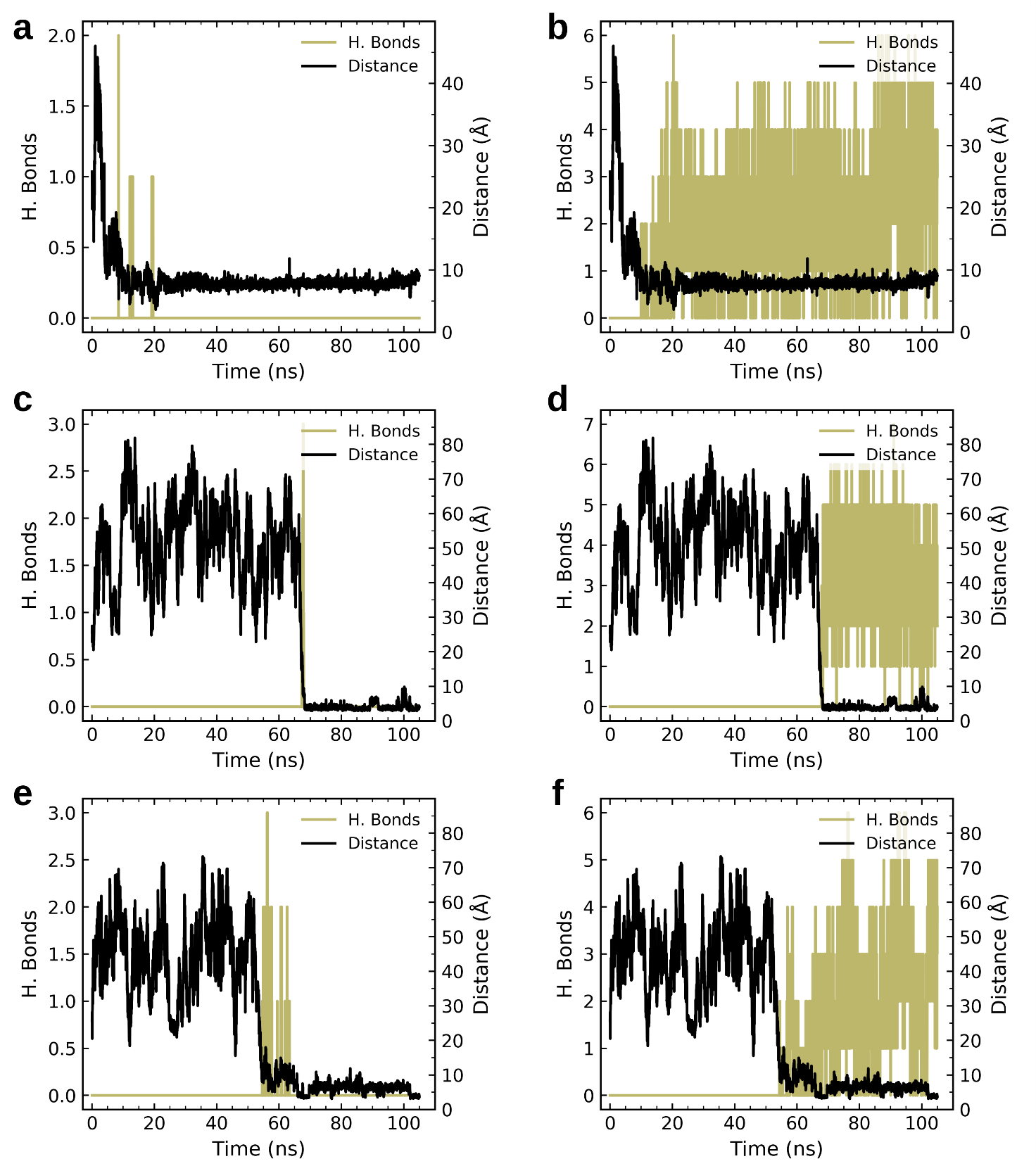

**Figure S7.** Traces of number of hydrogen bonds and distance between the glutamine carboxyl carbon and the guanidine carbon in Arg114. Two sets of residues are used for the analysis. Entry pocket residues set: Thr413 and Tyr194. Binding pocket residues set: Ser463, Asn411, Arg114, Asp433 and Thr409. Panels **a** and **b** shows the number of entry and pocket residue hydrogen bonds for R10, respectively. Panels **c** and **d** display the number of hydrogen bonds for R20 as explained above. Panels **e** and **f** show number of hydrogen bonds for R50 as described above.

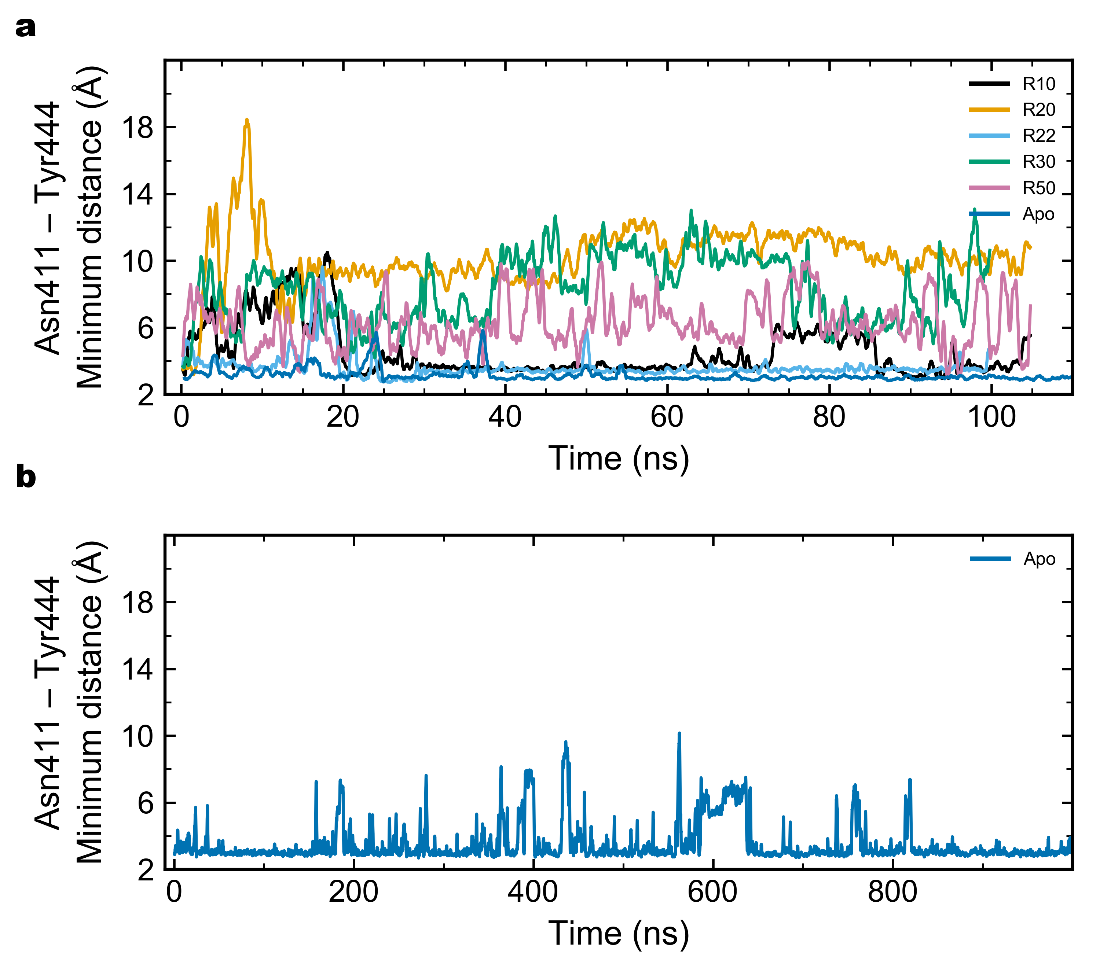

**Figure S8.** Minimum distance traces between heavy-atoms of Asn411 and Tyr444. **a,** aMD runs and conventional MD of apo EcoGGT. Stable receptor-substrate complexes are formed in trajectories R10, R20 and R50. R22 and R30 are trajectories where the substrate fails to bind. **b,** Extended conventional MD of apo EcoGGT. Data is smoothed by moving average over 20 frames (400 ps for aMD runs and 800 ps for conventional MD).

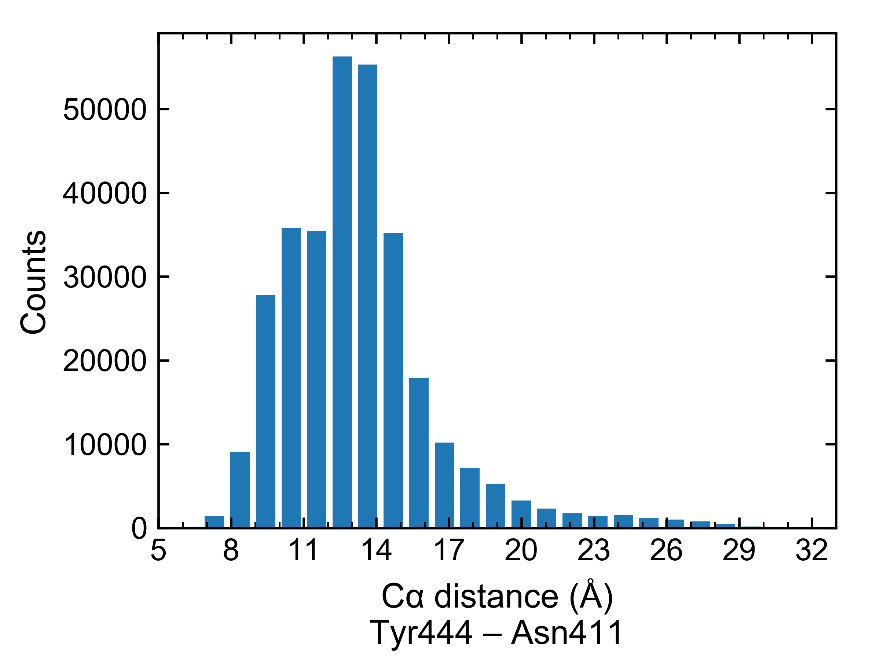

**Figure S9.** Histogram of number of biased aMD snapshots versus the Tyr444 – Asn411 Cα-distance. Number of bins is 25.

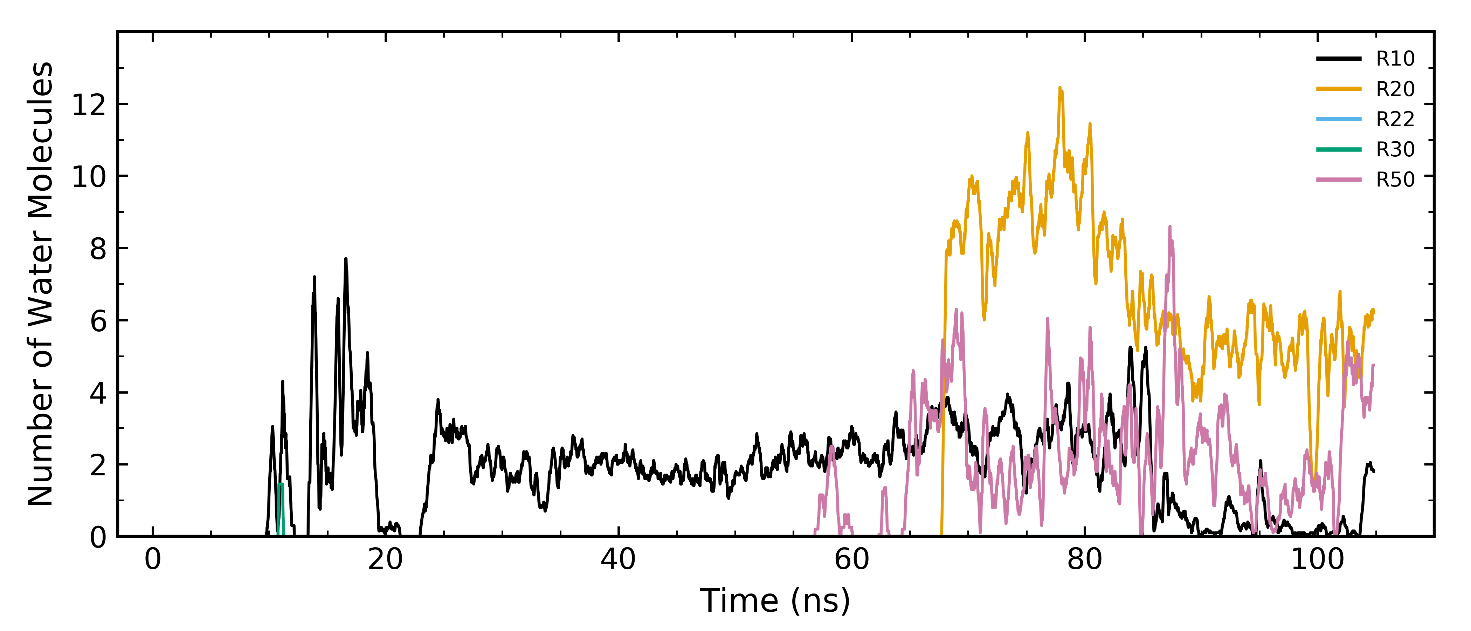

**Figure S10.** Number of water molecules inside a box containing glutamine and Tyr444 for substrate-binding aMD trajectories R10, R20 and R50. R22 and R30 are trajectories where the substrate fails to bind. The box is defined by the minimum and maximum coordinates of heavy-atoms atoms in glutamine and Tyr444 sidechain. Water molecules are counted when inside the box and within 7 Å of both glutamine and Tyr444. Calculation is carried out when glutamine’s carboxyl carbon is within a sphere of radius 8 Å centered at the guanidine carbon of Arg114. Water selection is updated for each frame. Data is smoothed by moving average over 20 frames (400 ps). The VMD script used for the calculations is included in the Supplementary Material.

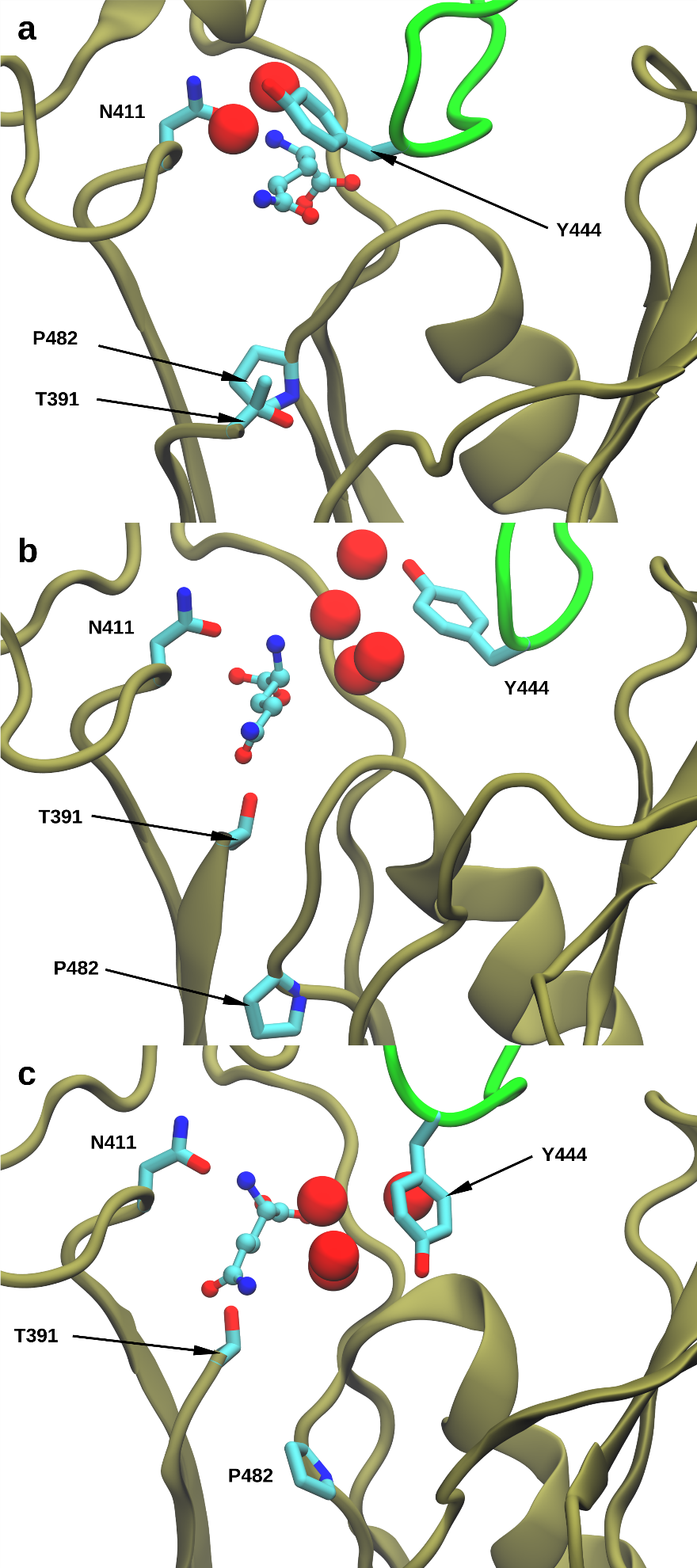

**Figure S11.** Glutamine-EcoGGT complexes corresponding to the last frames of the following aMD trajectories: a) R10. b) R20. c) R50. Glutamine represented in ball and sticks. Lid loop and chain B are shown in green and tan ribbons, respectively. Nitrogen, carbon and oxygen atoms are represented in blue, cyan and red color spheres, respectively. Oxygen atoms of water molecules are shown in red space filled representation.

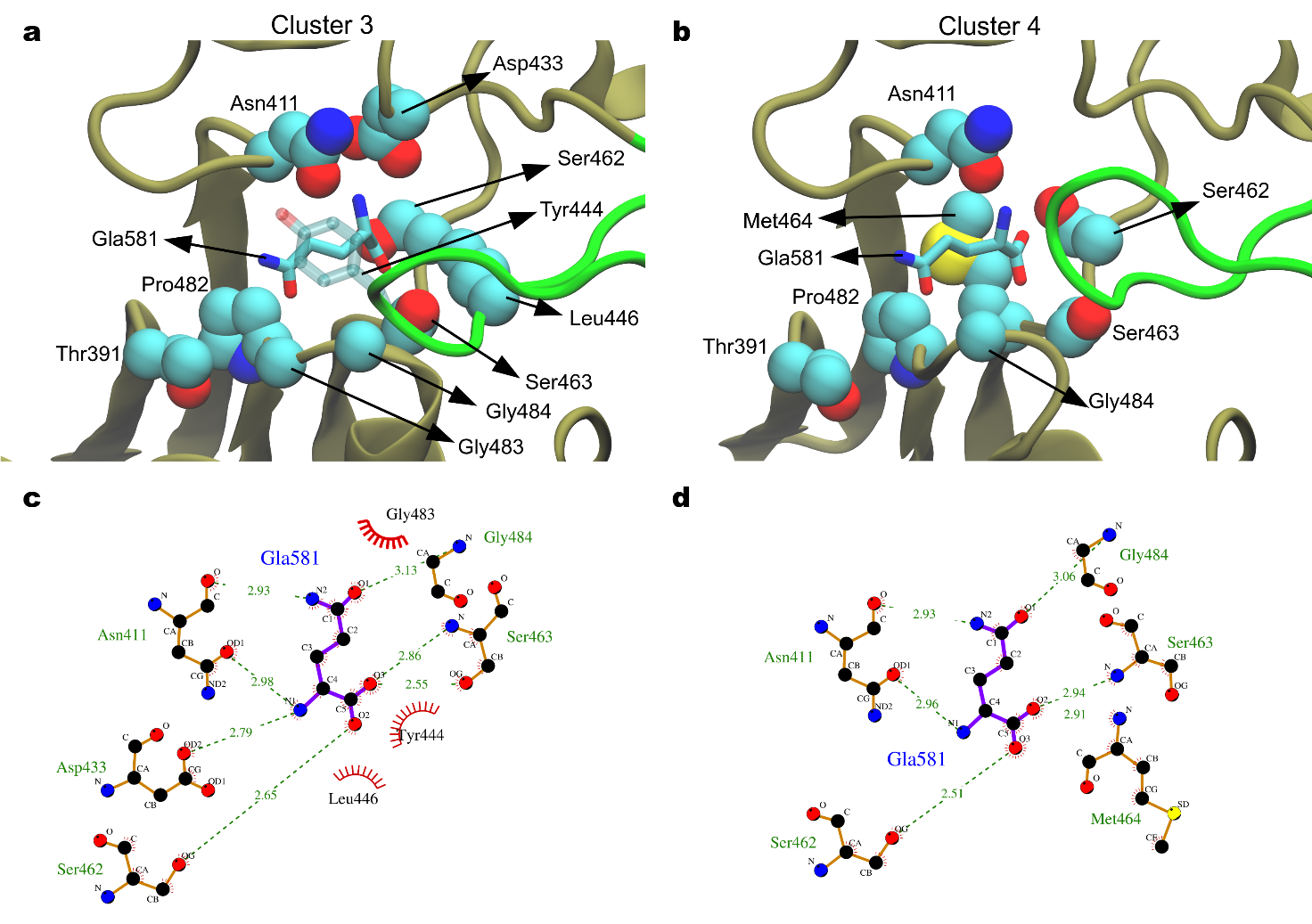

**Figure S12.** Structural representation of clusters 3 and 4. Cα distance between Tyr444 – Asn411 of 10 Å and 9 Å, respectively. **a,** Binding pocket of the representative structure of cluster 3. Tyr444 is shown in transparent licorice representation. **b,** Binding pocket of cluster 4. **c,** Hydrogen bond network and hydrophobic contacts of the glutamine (Gla 581), and the protein pocket in the representative structure of cluster 3. **d,** Receptor-substrate interactions plot of the representative structure of cluster 4. The substrate and protein residues involved in hydrogen bonding and hydrophobic interactions are highlighted in licorice and space-filling representation, respectively. Lid loop is shown in green ribbons. Only chain B is shown in tan ribbons. Nitrogen, carbon and oxygen atoms are represented in blue, cyan and red color spheres, respectively. Two-dimensional visualization obtained with Ligplot+.

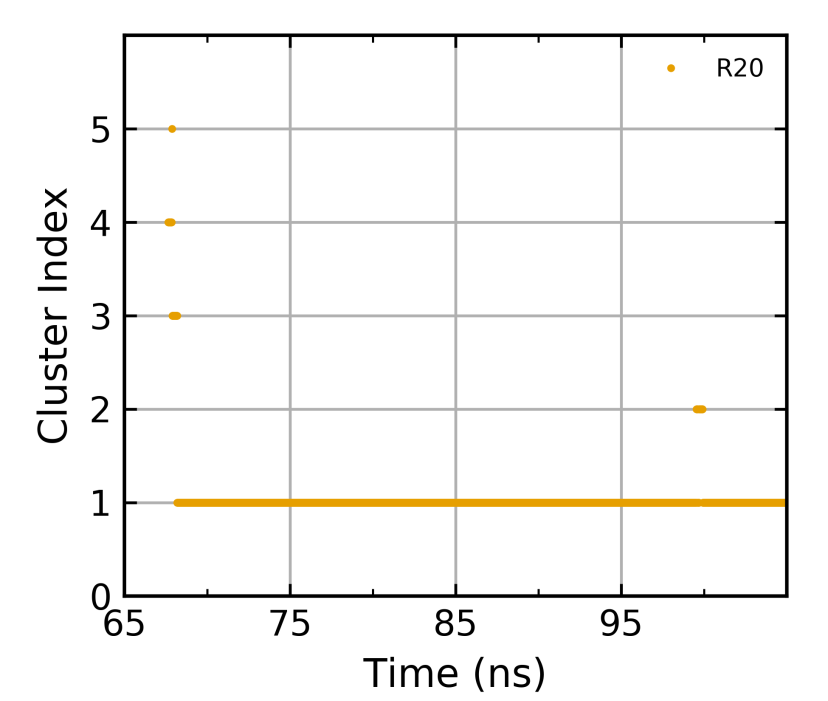

**Figure S13**. Plot of clusters versus simulation time for trajectory R20. The cluster conformations are reported in Figure 7.

VMD script to calculate number of waters between the substrate and the lid-loop:

\# Protein residue indexes in Amber Topology: 1 to 541

\# Tyr444 (resid 405)

set topol topol.prmtop

set traj_amber whole_run10.nc

\# distance cutoff for water contacts in Angstroms

set cutoff 7

\# distance cutoff between Arg114 (resid 78) and glutamine amide carbon in Angstroms

set distance 8

mol new $topol

mol addfile $traj_amber type netcdf waitfor all

set watgroup [atomselect top "(same fragment as water within $cutoff of resname GLA) and (same fragment as water within $cutoff of resid 405) and name O"]

set sel [atomselect top "(resname GLA and not hydrogen) or (resid 405 and sidechain and not hydrogen)"]

set amidecarbon [atomselect top "resname GLA and name C5"]

set hydroxyl [atomselect top "resid 78 and name CZ"]

set n [molinfo top get numframes]

set outfile [open "number_of_waters_${cutoff}A_d${distance}.dat" "w"]

puts -nonewline "\n Writing output..."

set step 1

for {set i 0} {$i < $n} {incr i $step} {

molinfo top set frame $i

$watgroup update

$sel update

$amidecarbon update

$hydroxyl update

\# vmd ouputs the result in a list

set mobile [lindex [$amidecarbon get {x y z }] 0]

set ref [lindex [$hydroxyl get {x y z }] 0]

set dist [veclength [vecsub $mobile $ref]]

set dims [measure minmax $sel]

set vecmin [lindex $dims 0]

set vecmax [lindex $dims 1]

set xmin [lindex $vecmin 0]

set ymin [lindex $vecmin 1]

set zmin [lindex $vecmin 2]

set xmax [lindex $vecmax 0]

set ymax [lindex $vecmax 1]

set zmax [lindex $vecmax 2]

set ids [$watgroup get resid]

set withinbox {}

set ninsidebox 0

set waternum [$watgroup num]

set coords [$watgroup get {x y z }]

foreach coord $coords idx $ids {

lassign $coord x y z

if { ($x > $xmin) && ($x < $xmax) && ($y > $ymin) && ($y < $ymax) && ($z > $zmin) && ($z < $zmax) } {

set ninsidebox [expr {$ninsidebox + 1}]

lappend withinbox $idx

}

}

if { $dist < $distance} {

puts $outfile "$i $waternum $ninsidebox All resid: $ids Within box resid: $withinbox"

} else {

puts $outfile "$i $waternum 0 All resid: $ids Within box resid: $withinbox LATERAL CONTACT"

}

}

close $outfile

puts -nonewline " Done !\n\n"

quit
